# Ultradeep N-glycoproteome Atlas of Mouse Reveals Spatiotemporal Signatures of Brain Aging and Neurodegenerative Diseases

**DOI:** 10.1101/2025.02.15.638397

**Authors:** Pan Fang, Xiangming Yu, MengYang Ding, Qifei Cong, Hongyu Jiang, Qi Shi, Weiwei Zhao, Weimin Zheng, Yingning Li, Zixiang Ling, Wei-Jun Kong, Pengyuan Yang, Huali Shen

**Author notes:** Correspondence: Huali Shen, Pan Fang. These authors contributed equally to the manuscript (Pan Fang, Xiangming Yu).

## Abstract

The current depth of site-specific N-glycoproteomics is insufficient to fully characterize glycosylation events in biological samples. Herein, we achieved an ultradeep and precision analysis of the N-glycoproteome of mouse tissues by integrating multiple workflows. The largest N-glycoproteomic dataset to date was established on mice, which contained 91,972 precursor glycopeptides, 62,216 glycoforms, 8,939 glycosites and 4,563 glycoproteins. The database consisted of 6.8 million glyco-spectra (containing oxonium ions), among which 160,928 were high-quality spectra with confident N-glycopeptide identifications. The large-scale and high-quality dataset enhanced the performance of current artificial intelligence models for glycopeptide tandem spectrum prediction. Using this ultradeep dataset, we observed tissue specific microheterogeneity and functional implications of protein glycosylation in mice. Furthermore, the region-resolved brain N-glycoproteomes for Alzheimer’s Diseases, Parkinson Disease and aging mice revealed the spatiotemporal signatures and distinct pathological functions of the N-glycoproteins. A comprehensive database resource of experimental N-glycoproteomic data from this study and previous literatures were further established. This N-glycoproteome atlas serves as a promising tool for revealing the role of protein glycosylation in biological systems.

## Introduction

Glycosylation is one of the most widespread and essential post-translational modifications of proteins, characterized by diverse, structurally complex, and dynamic glycan structures^1, 2^. Glycans significantly impact protein functions, playing crucial roles in physiological and pathological processes^3, 4^. Glycoproteins typically feature multiple glycans with different structures at individual sites within the polypeptide. N-glycoproteins are defined by glycans attached to specific asparagine residues within the consensus sequence N-X-S/T/C (X ≠ P). In vivo glycoconjugate synthesis is a dynamic process that depends on the local milieu of enzymes, sugar precursors and organelle structures as well as the cell types involved and cellular signals^5^. Identifying glycoproteins and characterizing the composition and structure of site-specific glycans are crucial for understanding the mechanisms underlying health and disease^6-8^. This underscores the necessity of experimentally determining N-glycosylation sites and glycans in vivo.

Mass spectrometry-based N-glycoproteomics faces several challenges including macro- and micro-heterogeneity, the low abundance of glycopeptides, incomplete enrichment for single method, and poor-quality and complex mass spectra caused low spectral identification rates and false discovery^9-15^. Significant efforts have been made to improve sample preparation, enrichment methods, spectral acquisition, and the development of various software for spectral interpretation^14, 16-25^. However, the identification depth and precision of N-glycoproteomics still need improvement.

In our previous study, we reported the largest mouse N-glycoproteome dataset at that time using pGlyco 2.0^19^. This high-quality dataset has supported the development of multiple glycosylation software such as MSFragger-Glyco, StrucGP, Glyco-Decipher, GlycanFinder and GlycReSoft^26-30^. Currently, artificial intelligence algorithms were employed for glycopeptide spectrum prediction, identification, and quantification, requiring large volumes of high-quality training data^31-33^. Therefore, there is an urgent need to establish an even larger foundational glycoproteomics dataset to meet these demands.

Here, we significantly expanded the mouse N-glycoproteome by implementing complimentary workflows across five different tissues, utilizing three enzymes, two enrichment methods, and performing five LC-MS/MS replicates (Figure 1). In total, we acquired 8,950,826 MS/MS spectra, of which 6,854,267 (76.6%) were glyco-spectra containing oxonium ions (Figure S1). From this data, approximately 1,041,225 glycopeptide spectra matches (GPSMs) were identified using multiple software. The confidence levels of these GPSMs categorized as high, moderate, low, and ambiguous-were determined based on the consistency of identifications across different software. This extensive analysis led to the identification of 91,972 unique precursor glycopeptides, 62,216 unique glycoforms, 8,939 glycosites, and 4,563 glycoproteins. This represents the largest multi-organ glycoproteomics dataset for mice to date, offering a more comprehensive and precise depiction of both the macro- and micro-heterogeneity of the mouse N-glycoproteome. We further conducted spatial and temporal glycoproteomic studies of brain to investigate the aging process and two neurodegenerative diseases, Alzheimer’s disease (AD) and Parkinson’s disease (PD), revealing the significant role of glycosylation in both aging and diseases, with distinct spatiotemporal characteristics. To facilitate access to this data, we developed a web-based tool, N-GlycoMiner (www.NGlycoMiner.com), which allows users to query site-specific glycan and tissue-specific glycoproteins. Combining the results of this study with existing literature, the platform currently hosts data on 310,416 glycoforms, 37,940 glycosites, and 12,078 glycoproteins for mice. This resource provides ultra-deep and precise analyses of experimental mouse N-glycoproteomes, serving as a valuable asset for the scientific community.

**Figure 1.**
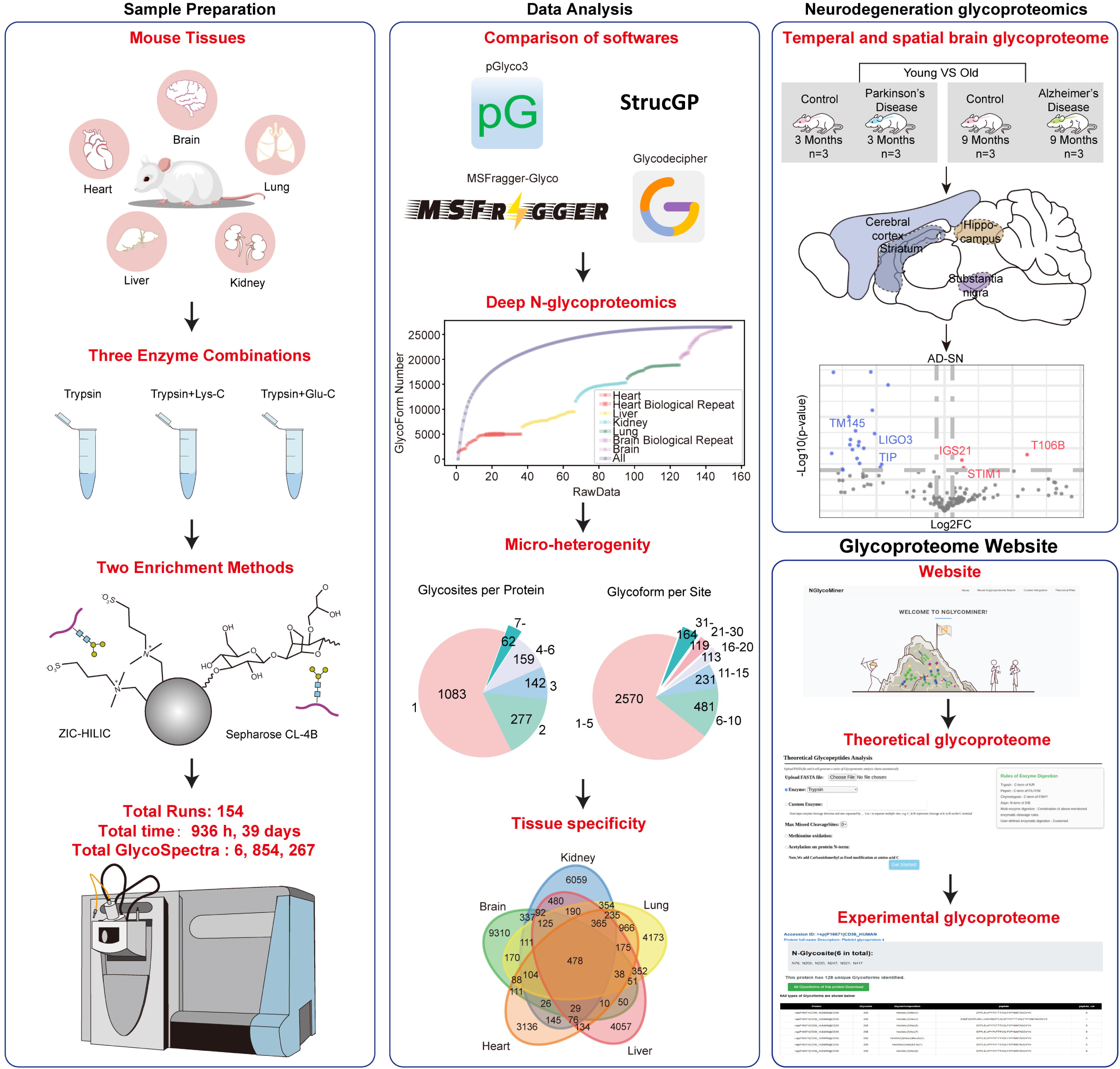
Comprehensive workflow of mouse N-glycoproteomics study, including sample preparation, data analysis, neurodegeneration glycoproteomics and N-glycoproteome website. Sample preparation: Five kinds of mouse tissues (brain, lung, kidney, liver and heart) were studied. Three enzyme combinations (Trypsin, Trypsin coupled with Lys-C and Trypsin coupled with Glu-C) were applied for protein digestion. Two enrichment methods (ZIC-HILIC and Sepharose CL-4B) were utilized. A total of 154 raws over 936 hours (39 days) were acquired with 6.8 million glyco-spectra (containing oxonium ions 204.0866). Data Analysis: Four software (pGlyco3, StrucGP, MSFragger-Glyco and Glyco-Decipher) were compared. Deep N-glycoproteomics analysis was conducted, showing the number of glycopeptides identified in different tissues. Micro-heterogeneity and tissue specificity was exhibited. Neurodegeneration Glycoproteomics: Temporal and spatial brain glycoproteome analyses were performed, comparing control, Parkinson’s disease, and Alzheimer’s disease states at different ages (3 and 9 months). Multiple brain regions, including hippocampus (HPC), prefrontal cortex (PFC), substantia nigra (SN), and striatum (STR), were analyzed. Glycoproteome Website: The results were available on a dedicated N-glycoproteome website, featuring experimental N-glycoproteome data.

## Results

### Ultradeep N-Glycoproteome Profiling of Mouse Tissues Using Complementary Methods

Based on the analysis of peptide sequences from the canonical mouse proteome FASTA file, 81% of the proteins contain the sequon N-X-S/T/C (where X ≠ P) and are theoretically likely to be N-glycosylated (as analyzed on our website below). The current depth of N-glycoproteomics identification was insufficient to describe the spatiotemporal specificity of tissues and cells during biological and pathological processes. In this study, we used complementary methods to perform the N-glycoproteome profiling of five mouse tissues, achieving extraordinary identification depth and precision.

The overview of experimental design was showed in figure 1. The ultradeep N-glycoproteome profiling of five mouse tissues, including brain, lung, kidney, liver and heart were performed using three kinds of enzyme combinations (trypsin, Lys-C coupled trypsin, and Glu-C coupled trypsin), two enrichment methods (ZIC-HILIC and Sepharose CL-4B) and five LC-MS/MS replicates with optimal LC (6 hours) and MS methods (Figure 1). This comprehensive approach resulted in a total of 154 runs (5 tissues × 3 enzymes × 2 enrichment methods × 5 replicates) conducted over 936 hours (excluding washing steps) across 39 days (Supplementary Data 1). This study achieved high reproducibility in both technical and biological replicates (Figure S2). The combination of different enzymes and enrichment methods significantly increased the depth of the N-glycoproteomics, and their underlying preferences were discussed in detail in Supplementary Note 1 (Figure S3 and Figure S4). By employing the above complementary methods, we achieved saturated glycoform identification counts (from pGlyco3 results) for each tissue and methods, demonstrating the robustness and completeness of the glycoproteomic analysis (Figure S5).

### The Largest Mouse N-Glycoproteome Database Enabled by Multi-Engine Integration and Confidence-Centric Data Curation

With the development of various glycoproteomics search engines^23^, evaluating and utilizing the identification results from different engines has become an indispensable issue. We compared four widely used and free available software, including pGlyco 3.0, StrucGP, Glyco-Decipher and MSFragger-Glyco. With 0.3-0.7 million GPSMs identified by each engine, approximately 1 million (1,041,225) GPSMs out of the total (nearly 9 million) were identified by the four search engines (Figure 2a). In comparison of the multi-level identifications, the ranking of identification numbers at the GPSM, precursor, and glycoform levels were almost consistent across all tools, with Glyco-decipher achieving the highest, followed by MSFragger-Glyco, pGlyco3, and finally StrucGP, which had the lowest (Figure 2a, Figure S6a, b). However, at glycosite, glycoprotein and glycan composition levels, the results vary significantly (Figure S6c, d, e). pGlyco3 showed the lowest level in glycosite and glycoprotein identifications but performs moderately well in other categories. StrucGP performed comparably to MSFragger-Glyco and Glycodecipher at the glycosite and glycoprotein levels. This analysis underscores the trade-offs between the tools in their ability to identify glycoforms, glycoproteins, glycosites and glycans, highlighting the influence of their design and analytical focus on identification performance. In comparison of the preferences or bias among these tools, we found that StrucGP consistently identified shorter peptides and glycans, whereas MSFragger-Glyco shows a preference for longer peptides, and Glyco-decipher tends to identify longer glycans (Figure S7a, b, c). In terms of glycan types (Figure S7a), StrucGP is biased toward high-mannose and pauci-mannose glycans. MSFragger-Glyco exhibits the highest sialic acid content in its identifications. Both pGlyco3 and Glyco-decipher demonstrate a stronger focus on fucosylated glycan identifications. The analysis of software tools for quantifying glycoproteomics data in heart and brain samples revealed high consistency in glycoprotein and glycosite quantification, but low consistency at the glycan and site-specific glycoform levels. Correlation analysis showed strong agreement in glycoprotein and glycosite quantification (Pearson coefficients >0.78), but lower agreement for glycans (Pearson coefficients 0.52-0.67). These findings highlight the need for caution when comparing glycan and glycoform data across different tools, underscoring the importance of improving glycan quantification algorithms and validating using other experiments (Figure S8). The detailed comparison of the four softwares were presented in Supplementary Note 2.

**Figure 2.**
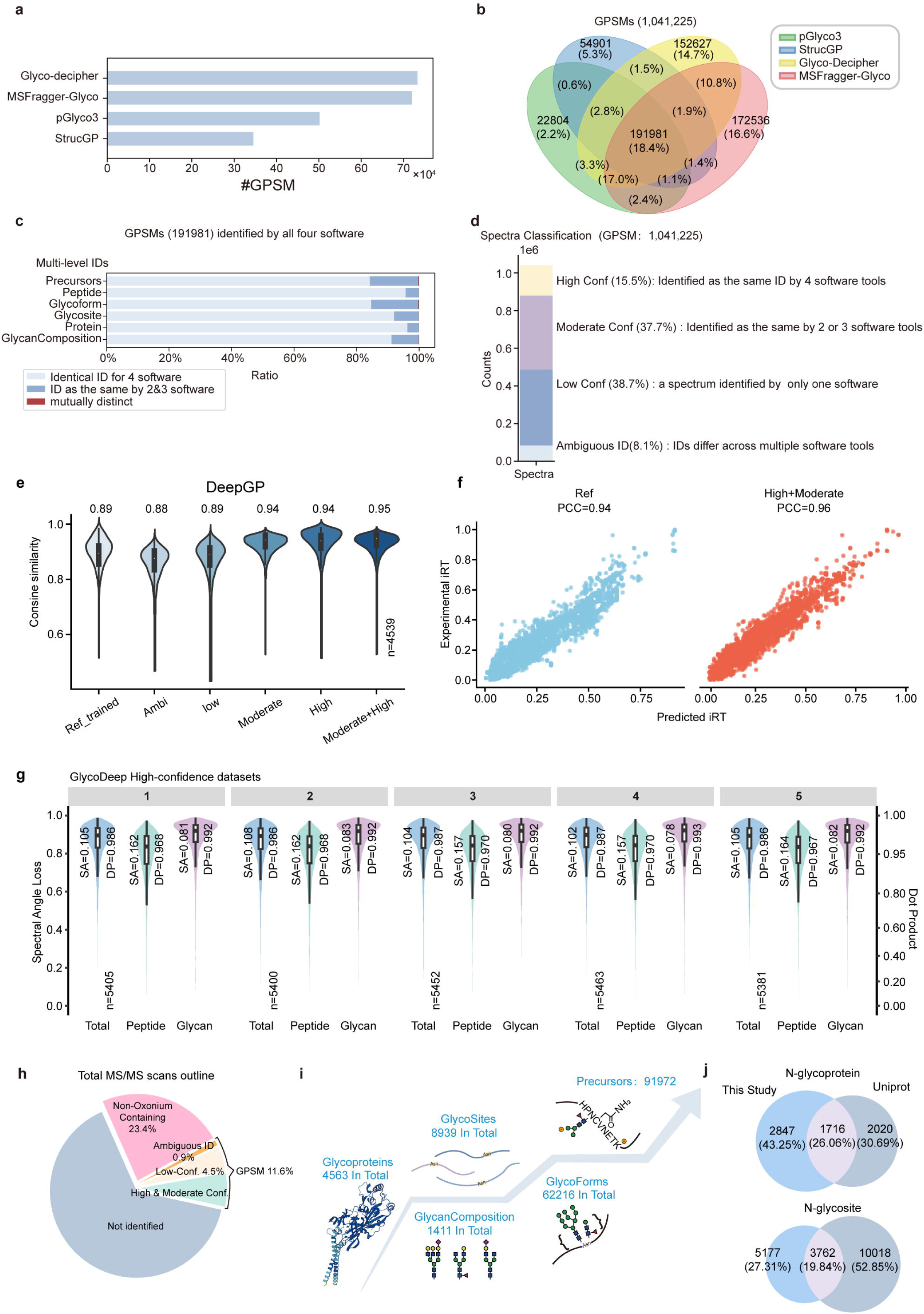
Comprehensive analysis of identifications from multiple software. (a) The number of GPSMs identified by four software. (b) A Venn diagram illustrating the overlap of GPSMs identified by the four software, highlighting unique and shared identifications among them. (c) The common identified GPSMs by the four software can result in consistent (Identical ID for 4 software) and inconsistent identifications (ID as the same by 2 or 3 software, ID mutually distinct) in precursors and other identifications. The ratios of consistent and inconsistent identifications were showed. (d) Spectra classification based on confidence levels. (e) Distribution of cosine similarities between the predicted and experimental spectra for different training datasets using DeepGP model: literature-reported models, ambiguous, low, moderate, high_split, and high_split+moderate subsets. The remaining high-confidence subset (4539 spectra) were utilized for testing. (f) Correlation between the predicted and experimental iRT is presented, with Pearson correlation coefficients (PCC) shown above. Predictions were made using model provided in the literature and the model trained on high and moderate-confidence data, without RT calibration. (g) Distribution of spectral similarities for high-confidence dataset in 5-fold cross validation using DeepGlyco model. (h) Pie chart outlining the total MS/MS scans, categorized by confidence levels and types of spectral identifications. (i) Overview of glycoproteome data, showing the total number of glycoproteins, glycosites, and glycoforms identified in the study. (j) Comparison of data from this study with that from UniProt database entries. In (e) and (g), the median values of cosine similarities, spectral angle loss (SA) and dot product (DP), as well as data size (n) are indicated respectively. The lower and upper hinges of the box represent the first and third quartiles. The lower and upper whiskers extend from the hinges to the smallest and largest values within 1.5 times the interquartile range (IQR). Source data are provided as a Source Data file.

We then evaluated the identification confidence of all the results. As the Venn diagrams of GPSMs showed in Figure 2b, 191,981 GPSMs were identified by all of the four search engines, 160,928 of which were identified as the same glycopeptide precursors. These 160,928 GPSMs and the corresponding glycopeptide precursors were considered highly confident identifications. Although each search engine has undergone strict quality control, including scoring system and FDR, there were still a number of cases where multiple software identified inconsistent results for the same spectrum (Figure 2c). We analyzed the distributions of the inconsistent identifications from four or three software (Figure S9). The analysis of identification inconsistencies among the four software tools revealed that pGlyco3 exhibited the highest reliability, while MSFragger-Glyco identified more spectra but with greater inconsistency, highlighting a trade-off between sensitivity and accuracy. We thus assumed that the consistent results from more than one software have higher confidence and proposed a classification of all the identified GPSMs based on the confidence of them (Figure 2d): high confidence spectra (a spectrum was identified as the same precursor glycopeptide by all the four software) (number160,928, 15.5%), moderate confidence spectra (a spectrum was identified as the same precursor glycopeptides by at least two of the four software) (number 392,030, 37.7%), low confidence spectra (a spectrum was only identified by one software) (number 402,705, 38.7%) and ambiguous spectra (a spectrum was identified by different software as totally different precursors) (Supplementary Data 2) (84,585, 8.1%).

The high-quality glycopeptide spectra with their high-confidence identification results, could make up for the current lack of glycopeptide standard libraries and provide reliable, high-quality training data for AI model training (Supplementary Data 3). Here, we showed the examples of using the different confidence datasets produced in this study to retrain the current available DeepGP and DeepGlyco models^32, 33^. We trained DeepGP with different confidential datasets, including 90% of high-confidence data (High_train), moderate, low, ambiguous and a combination of moderate and high_train (Moderate+High-train), with remaining 10% of high-confidence data reserved for testing (Figure 2e). The results demonstrated that data with moderate confidence and above (High_train and Moderate+High-train) significantly improved the cosine similarity between the predicted and experimental mass spectra, with the median for Moderate+High_train reaching 0.95, outperforming the previously reported model DeepGP. The moderate-confidence dataset surpassed the high-confidence dataset, likely due to the larger size of the moderate dataset (60k vs. 26k glycopeptides) and the relatively high data quality, as it contains spectra co-identified by two or three different search engines. Also, the model trained on Moderate+High_train data achieved higher Pearson correlation coefficient (PCC) of iRT prediction than the previously reported model in the absence of RT calibration (Figure 2f). In the other DeepGlyco model, we performed 5-fold cross-validation on the high-confidence dataset, where four parts were used for training and one part for validation in each iteration (Figure 2g). The median dot product (DP) for the whole glycopeptides was consistently above 0.986, and exceeded 0.992 for the glycan parts across all subsets. Furthermore, the spectral angle loss (SA) for glycopeptides ranged from 0.102 to 0.108, while for the glycan parts, it ranged from 0.078 to 0.083. The results were stable, indicating superior data quality and representativeness (Figure S10). These tests on current AI models demonstrated that the data with different confidence can facilitate MS/MS or retention time prediction and developing AI-driven software. (detailed discussion in Supplementary Note 3).

Analysis of the total ms/ms scans showed that 23.3% of total spectra did not contain the oxonium ions, and 65.1% of total spectra were not identified by any software (Figure 2h). For each glycoproteomics software, the spectra interpretation rate (GPSM/total spectra) ranges from 3.8%-8.2%, which is surprisingly low. Even with the combination of the four software, which has greatly increased the identification depth of glycoproteomics, the spectra interpretation rates only increased to 11.6%. There were still a large number of un-identified glyco-spectra (88.4% of total spectra) and the low-confidence and ambiguous identifications (46.8% of GPSMs) (Figure S11). The improvement in mass spectrometry techniques, software identification, and AI-based spectral recognition is still urgent.

After removing the ambiguous identifications, a total of 956,640 GPSMs were retained, from which 91,972 unique precursor glycopeptides and 62,216 unique glycoforms were identified (Figure 2i). Then 8,939 glycosites (protein and site) with 1,411 glycan compositions on 4,563 glycoproteins were derived. This study provided more glycoproteins compared to the glycoproteins in Uniprot (3736), with 2847 proteins and 5177 glycosites additionally identified (Figure 2j). It greatly expanded the scale of moue N-glycoproteome database we built in 2017^19^. The numbers of GPSMs and glycoforms in this study were 13 times and 6.2 times of that from 2017, respectively (Figure S12a). To our knowledge, this is the largest dataset for mouse N-glycoproteome to data.

We compared the identification numbers of five tissues, three enzyme combinations and two enrichment methods (Figure S12 b-d, Figure S13). About 16,000-29,000 glycopeptide precursors, 11,000-20,000 glycoforms, 2,000-3,600 glycosites, and 1,100-2,000 glycoproteins were identified from different mouse tissues. The numbers of identifications in brain were the highest while those in heart were the lowest. A total of 62,216 glycoforms were identified in five tissues and only 671 (1.08%) glycoforms were overlapped in five tissues while up to 6107 glycoforms were exclusive to brain.

### Tissue Specific Micro-Heterogeneity of Mouse N-Glycosylation Modifications

To investigate the micro-heterogeneity and tissue specificity of glycosylation modifications, we utilized the identification and quantification results from pGlyco3, given its lower rate of inconsistent identifications and stringent quality control at both the peptide and glycan levels compared to other software (Figure S9, Supplementary Note 2 and Supplementary Data 4). The overall mass distribution of the intact glycopeptides, de-glycopeptides and glycans were exhibited, ranging from 2,000-6,000 Da for intact glycopeptides, 500-4,000 Da for de-glycopeptides and 1,000 - 3,000 Da for glycans (Figure 3 a, b and c, Figure S14a-e). The portion of glycan mass on a glycopeptide was from 19%-85% with median ratio as 55% (Figure S14a). The experimental de-glycopeptide mass distributions were showed in contrast to the theoretical ones (peptides with N-X-S/T/C (X ≠ P)). About 70% of the experimental de-glycopeptides mass were between 1,000-2,500 Da. The experimental and theoretical deglycopeptides mass distribution reflected the limitations of the mass range identifiable by mass spectrometry (Figure 3b, Figure S14c). In addition, we surprisingly observed that the brain glycans were generally smaller than that from the other tissues, which indicates a unique glycosylation profile in brain tissues compared to other tissues (Figure 3c). We further analysed the secondary structures of the N-glycoproteins using the AlphaFold2 database (Figure 3d). The results revealed that coil and loop are the most prevalent secondary structures, followed by β-strands and hydrogen-bonded turns. We quantified the frequency of co-occurrence of glycan pairs at a same site (Figure S14f). The size of the dots indicated the incidence of co-occurrence. High mannose and fucosylated glycans exhibited a pronounced tendency to co-occur with other glycan types.

**Figure 3.**
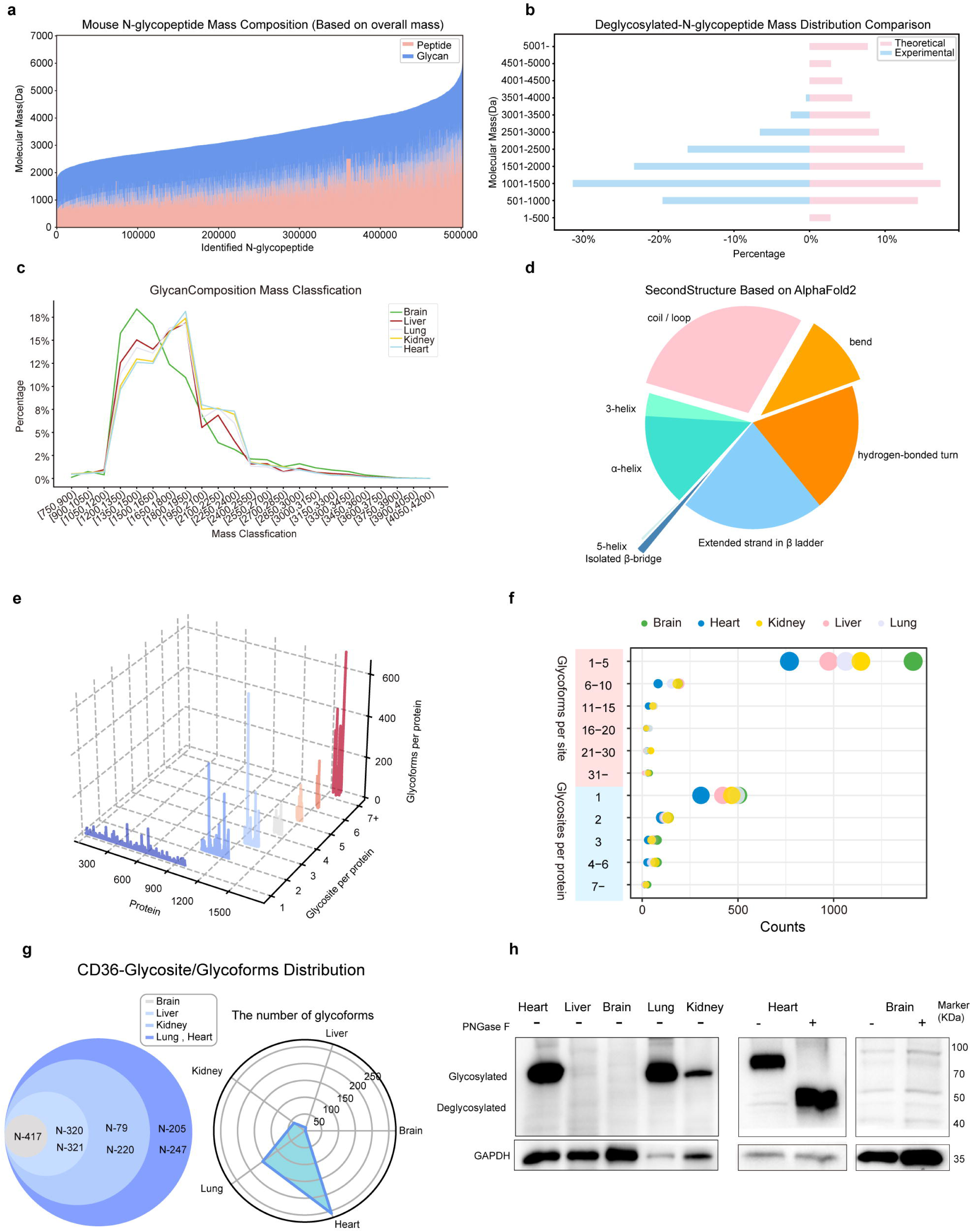
Comprehensive analysis of mouse N-glycoproteome. (a) Distribution of molecular masses for identified N-glycopeptides, illustrating the contributions from peptide and glycan components. (b) Comparison of theoretical and experimental molecular mass distributions of de-glycosylated N-glycopeptides. (c) Distribution of glycan composition mass across different mouse tissues. (d) Pie chart showing the distribution of secondary structure elements (coil, bend, hydrogen-bonded turn, extended strand in β ladder, 3-helix, α-helix, 5-helix, β-bridge) as predicted by AlphaFold2 and annotated by DSSP. (e) Three-dimensional histogram illustrating the distribution of glycoproteins by the number of glycosites and glycoforms per protein. (f) Scatter plot showing the distribution of glycosites and glycoforms per protein across different tissues. (g) Venn diagram and radar chart showing the distribution of glycosites and glycoforms of the CD36 protein across different tissues. (h) Western blot analysis of CD36 glycosylation in mouse tissues. Samples from heart, liver, brain, lung, and kidney were analyzed with (+) or without (-) PNGase F treatment. The experiments were repeated in four technical replicates with similar results.

The macro- and micro-heterogeneity of N-glycosylation from multiple tissues were comprehensively compared by analyzing the distribution of N-glycosites and site-specific N-glycans on proteins. We constructed a three-dimensional bubble plot (3D-hetero) that displays the number of N-glycosites (y axis) and the number of glycoforms (z axis) on each protein (x axis) (Figure 3e, Figure S14g-k). From a protein level, the huge heterogeneity on proteins with multiple sites (e.g. above 7) and up to 711 glycoforms on the proteins were revealed (AT1B1_MOUSE) (Figure S14l, m). On average, there are 2 sites on one protein and 7 glycans on one site in this dataset. This deep N-glycoproteome dataset exhibited much higher heterogeneity of site-specific glycopeptides than the previous studies^34^. The N-glycosylation heterogeneity for the five tissues were also analyzed (Figure 3f). The distributions of the number of glycosites and the number of glycoforms on proteins were compared among these tissues. Interestingly, the brain exhibited the highest micro-heterogeneity and the heart tissues exhibited the lowest micro-heterogeneity.

The tissue-specific glycoprotein expression was shown taking CD36 (Platelet glycoprotein 4) as an example. In the brain, only a single glycosylation site at N417 and 2 glycoforms of CD36 were detected (Figure 3g). However, in the heart and lungs, seven glycosylation sites were identified on CD36, with the heart notably having as many as 258 glycoforms (Supplementary Data 5). Xia ban. To validate the results from glycoproteomics, we performed additional Western blot analysis of CD36 (Figure 3h). The protein expression of glycosylated CD36 in the heart and lung was much higher than that in the brain. After removing the N-glycans with PNGase F, the N-glycans were completely removed, confirming that CD36 mainly undergoes N-glycosylation. In both conditions, we observed that the protein expression of glycosylated CD36 in the heart was much higher than in the brain. These results from the western blot analysis were consistent with the identification of glycoforms for tissues. The differences in glycosylated CD36 between heart and brain mainly stemmed from the protein expression levels in these two tissue types, which were higher in the heart and lower in the brain. CD36 has been documented to facilitate myocardial and adipose tissue fatty acid uptake in humans and recent study has reveal its role in promoting lung adenocarcinoma^35, 36^. The distinct differences of glycosylation on CD36 implied the importance of glycosylation for the tissue specific functions of CD36.

### Tissue-Specific Glycosylation Profiles and Functional Implications Revealed by Comprehensive Glycoproteomics Analysis

Furthermore, we comprehensively analyzed the tissue-specific N-glycosylation profiles in the five mouse tissues (Supplementary Data 4). Principle component analysis (PCA) of the glycoforms, glycans, glycosites and glycoproteins from different tissues showed that the N-glycosylation of brain was distinct from all the other tissues (Figure 4a and b, Figure S16a and b). Surprisingly, we found that kidney also presented a unique N-glycan pattern comparing with others (Figure 4a). The site-specific glycoforms in different tissues are weakly correlated, with the Pearson correlation values about 0.3-0.43 (Figure S16c).

**Figure 4.**
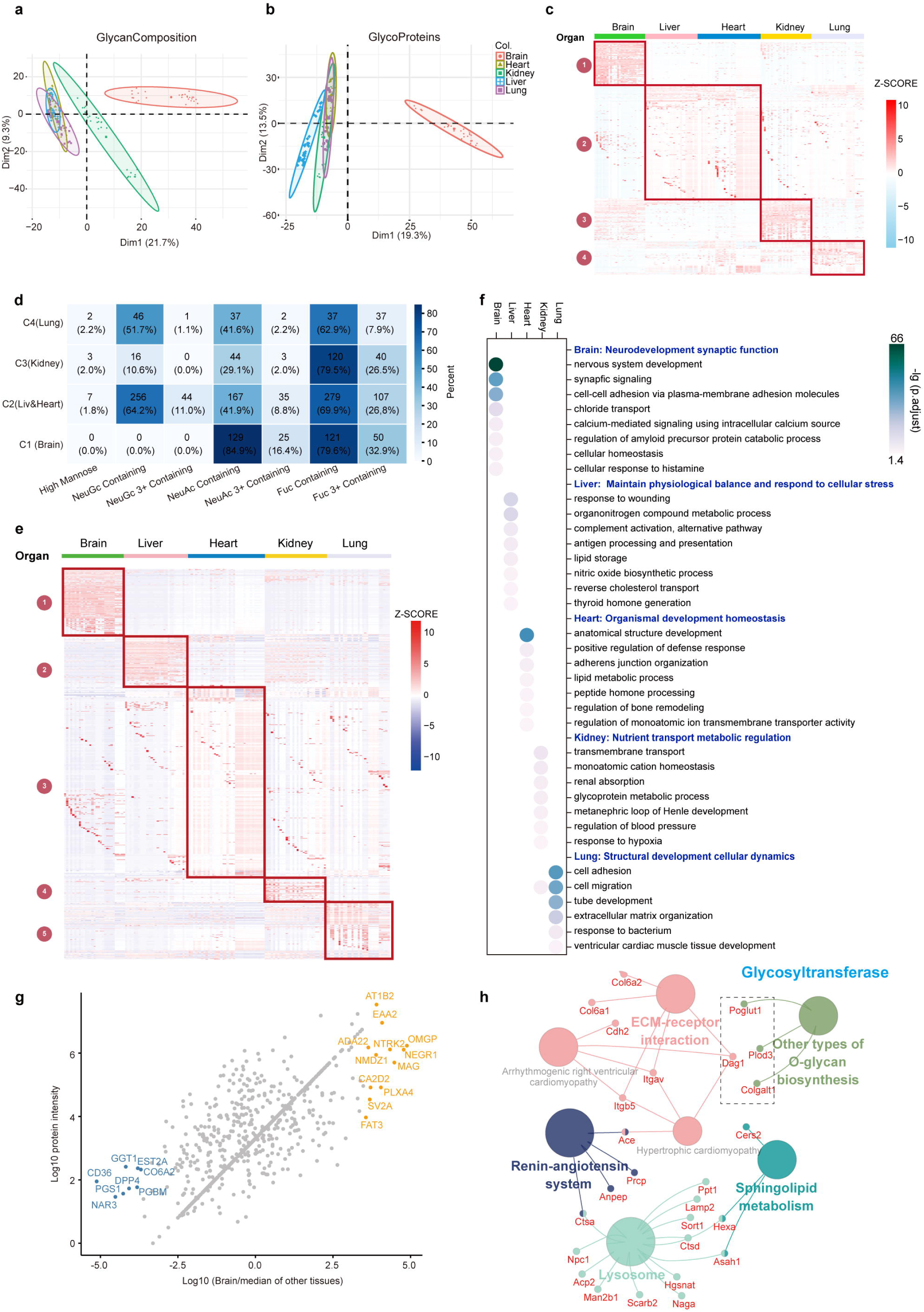
Comprehensive analysis of glycan composition and glycoproteins across mouse tissues. (a, b) PCA of glycan composition and glycoproteins across different mouse tissues. (c) heatmap showing hierarchical clustering of glycans across different mouse tissues, with z-scores indicating relative abundance. key clusters are highlighted. (d) heatmap showing the distribution of different glycan classes (High mannose, NeuGc containing, NeuGc 3+ containing, NeuAc containing, NeuAc 3+ containing, Fuc containing) in various tissues. (e) heatmap showing detailed glycoprotein expression levels across different tissues, with clusters highlighted for specific glycoprotein groups. (f) functional enrichment analysis showing the enriched biological processes for glycoproteins in different tissues. colors represent different tissues and dot size indicates the level of enrichment. P-values were calculated using a hypergeometric test and adjusted for multiple testing using the Benjamini–Hochberg FDR method. Terms with adjusted p-values < 0.05 were considered significant. (g) Scatter plot comparing the abundance of glycoproteins in the brain against the median abundance in other tissues. notable glycoproteins are highlighted. (h) glycoprotein interaction network analysis illustrating the interaction between glycoproteins involved in various biological processes and pathways.

Unsupervised clustering analysis of the N-glycans (Figure 4c and d) and N-glycoproteins (Figure 4e and f) expression were performed. Four distinct glycan subgroups (cluster 1-4) were characterized by tissue-specific features (Figure 4c). The glycan species of each cluster were further classified into glycan subtypes (Figure 4d). The glycan clusters highly expressed in brain tissue (cluster 1) primarily include fucose (79.6%) and NeuAc (84.9%), with no presence of NeuGc. Kidney specific glycan cluster (cluster 3) contain fucose (79.5%) and relatively low sialic acids. The glycans in liver and heart were collectively enriched in cluster 2 with high NeuGc (64.2%)

Five distinct glycoprotein clusters were characterized by tissue-specific features (Figure 4e). Next, we analyzed the biological processes associated with these tissue-specific glycoprotein clusters (Figure 4f). Glycoproteins enriched in the brain are primarily involved in neurodevelopment and synaptic function, including processes such as nervous system development, synaptic signaling, and cell-cell adhesion. Liver-specific glycoproteins mainly function in metabolism and cellular stress response. Heart-specific glycoproteins are involved in organismal development and homeostasis. Kidney-specific glycoproteins focus on transport and metabolic regulation, while lung-specific glycoproteins are associated with structural development and cellular dynamics.

We then illustrated the tissue-enriched glycoproteins in figure 4g. Taking brain as an example, the proteins, including AT1B2_Mouse (Sodium/potassium-transporting ATPase subunit beta-2), EAA2_Mouse (Excitatory amino acid transporter 2) et al, were glycosylated with higher abundance compared with other tissues. Another group of proteins (including CD 36, NAR3, PGS1, DPP4, et al) were less glycosylated in brain than other tissues. The specific glycoproteins in the other tissues were demonstrated in figure S16d-g.

Network analysis of the co-regulated N-glycopeptides/glycoproteins among the five tissues showed common processes regulated by N-glycosylation (Figure 4h). The co-regulated pathways included the other type of O-glycan biosynthesis, ECM-receptor interaction, sphingolipid metabolism, lysosome and renin-angiotensin system. Notably, the most prominently co-expressed hubgene, beta-hexosaminidase subunit alpha (Hexa), were mainly enriched in liver and lung tissues (Figure S16h). The genes marked in red were known protein-protein interactions. The high connectivity of Hexa with other N-glycoproteins underscores its role as a potential master regulator in both glycan synthesis and degradation.

### Spatial-temporal and Disease-specific N-glycoproteome in Mouse Brain

In the analysis of the tissue-specific glycoproteome, we found the brain glycoproteome is unique to all other tissues^37, 38^. To further elucidate the roles of glycosylation in brain functions, we performed a comparative N-glycoproteomic analysis on four mouse groups: Alzheimer’s Disease (AD, APP/PS1, 9 months), aging (wild type, 9 months), Parkinson’s Disease (PD, MPTP, 3 months), and young controls (wild type, 3 months), using samples from four brain regions: hippocampus (Hipp), prefrontal cortex (PFC), substantia nigra (SN), and striatum (Str) (Figure 1, Supplementary Data 1 and Supplementary Data 6).

The principal component analysis (PCA) revealed clear age-related separations between young and aging groups along the first dimension, while disease states (AD, PD) displayed less variation within the same age groups, indicating that age has a significant impact on the main variances (Figure 5a). Additionally, distinct spatial differences in protein glycosylation patterns were observed across the four brain regions. We identified significant disease- and age-specific glycosylation changes in various brain regions, as highlighted by volcano plots and illustrated in the upset plot (Figure 5b, Figure S17, S18). The upset plot illustrated the up-regulated and down-regulated glycoproteins in different conditions as well as the intersections of these conditions. Multiple glycoproteins, represented by the red bars, exhibited consistent changes across different brain regions under PD and aging conditions. The N-glycoproteomes exhibited significant temporal-, spatial-, and disease-specificity, as shown by the clustered heatmap (Figure 5c). This study found that aging mice (9 months) showed a significant decline in glycosylation levels of many proteins compared to younger mice (3 months), regardless of their health status (Figure 5c and 5d). This decline suggests compromised neural health, indicating that reduced glycosylation levels may serve as a potential biomarker for aging.

**Figure 5.**
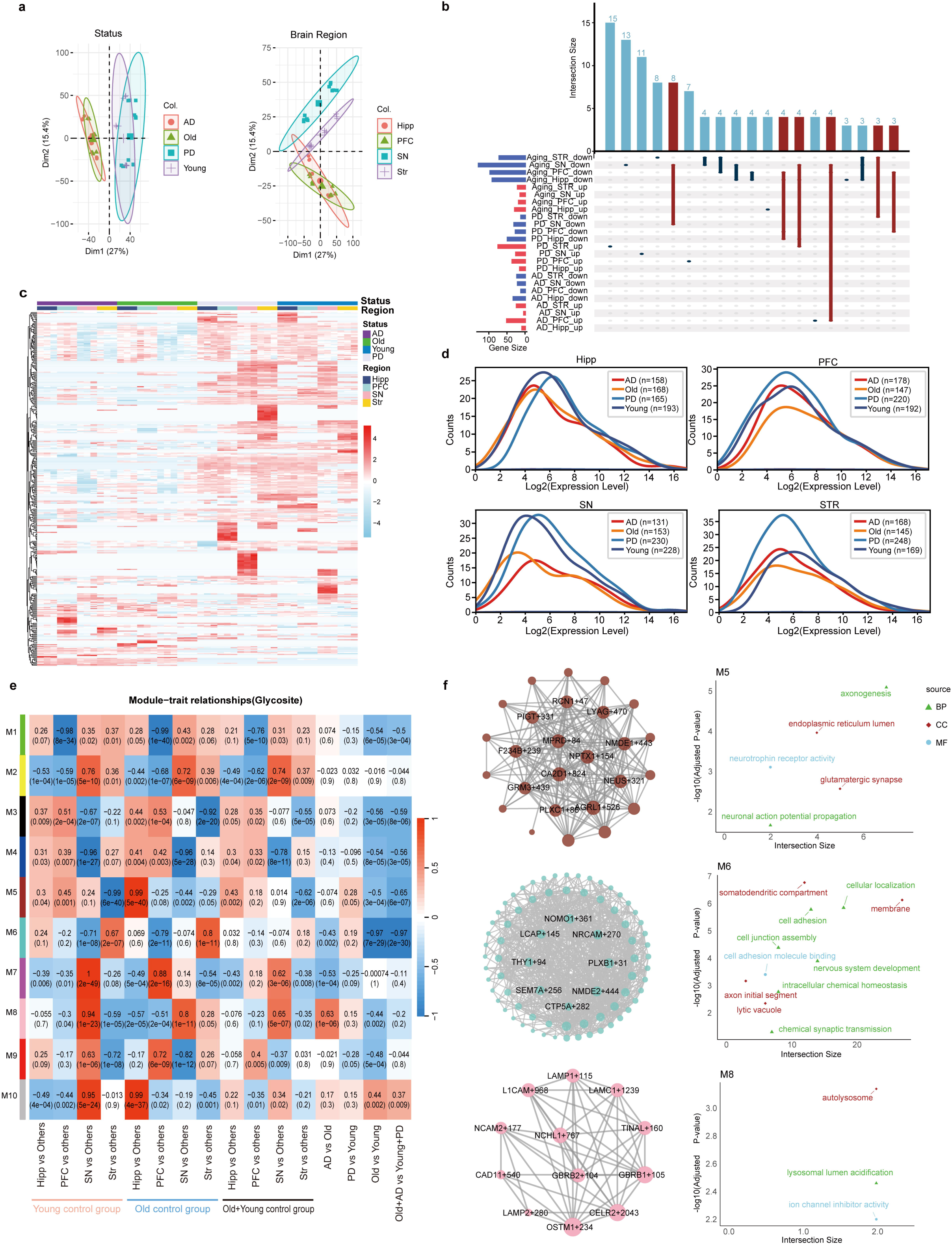
Comparative analysis of N-glycoproteomics in brain regions across different diseases and aging conditions. (a) PCA plots showing the clustering of samples based on glycoprotein profiles. (b) UpSet plot representing the intersection size of differentially expressed glycopeptides (DEGs) across various conditions and brain regions. The bars at the top indicate the number of DEGs unique to or shared between the conditions and regions. The left bar plot shows the overall gene set size for each condition/region combination. (c) Heatmap of glycopeptide profiles for key DEGs identified in the analysis. The rows represent glycopeptides, and the columns represent samples grouped by status and brain region. The color scale indicates the expression levels, with red representing high expression and blue representing low expression. (d) Density plots showing the distribution of log2 expression levels for different conditions across four brain regions. Each plot compares the expression distributions for AD, PD, old, and young conditions. (e) Module-trait relationship heatmap illustrating the correlation between glycosite modules (M1 to M10) and various traits, including status and brain region combinations. The color intensity indicates the strength of the correlation, with red representing positive correlations and blue representing negative correlations. The numbers in the cells represent the correlation coefficient and the p-value (in parentheses). (f) Network analysis of significant glycosite modules M5, M6 and M8. The left side shows the glycopeptides networks within each module, with hub glycosites highlighted. The right side displays the enriched Gene Ontology (GO) terms for each module, categorized into Biological Process (BP), Cellular Component (CC), and Molecular Function (MF). The green text indicates GO terms significantly associated with the respective modules. P-values were calculated using a hypergeometric test and adjusted for multiple testing using the Benjamini–Hochberg FDR method. Terms with adjusted p-values < 0.05 were considered significant. Source data are provided as a Source Data file.

To gain systems-level insights into glycan modifications in the brain and their changes across different regions and diseases, we constructed co-regulation networks of N-glycosites (Figure 5e and f) and N-glycans (Figure S19) by using the weighted gene co-expression network analysis (WGCNA)^39^. WGCNA clusters highly co-expressed N-glycosites or N-glycans into distinct modules, which facilitates identifying key modules and their associations with specific brain regions or disease conditions. In the heatmap, each cell displays the correlation coefficient and p-value. A total of ten modules were identified from the N-glycosite data. Notably, module M5 showed a strong negative correlation with the trait "Young Str vs. others" (r = -0.99, p < 6e-40), indicating significant downregulation in the substantia nigra of young subjects. M5 was also significantly enriched in the hippocampus of older mice and upregulated in Parkinson’s disease (PD). This module consisted of 26 co-regulated glycosites on 25 glycoproteins, with NPTX1_154, MPRD_84, and CA2D1_824 identified as hub genes. Gene Ontology (GO) enrichment analysis revealed that M5 is associated with biological processes such as axonogenesis and glutamatergic synapse function. Another module, M6, was enriched in the striatum in both young (r = -0.67, p < 2e-7) and old mice (r = -0.8, p < 1e-11), and it was significantly downregulated in Alzheimer’s disease (AD) (r = -0.43, p < 0.002) and during aging (Old vs. Young, r = -0.97, p < 7e-29). This module comprised 77 co-regulated glycosites on 65 glycoproteins, with THY1_94 and NRCAM_270 among the hub genes. GO terms for M6 include nervous system development and cell adhesion. Another noteworthy module, M8, was enriched in the substantia nigra in both young (r = -0.94, p < 1e-23) and old mice (r = -0.8, p < 1e-11), and significantly upregulated in AD (r = 0.63, p < 1e-6). M8 contained 12 co-regulated glycosites on 12 glycoproteins, centered around NPTN_283, and was highly involved in lysosomal functions. Details of the other modules are provided in the supplementary information (Figure S20).

The glycan modification co-regulation network analysis uncovered glycan modules associated with brain region specificity and disease pathology (Figure S19). For instance, the MEyellow (G1) module comprises 16 (80%) co-regulated, highly sialylated glycans that are enriched in the hippocampus of young subjects (Young_Hipp) and the striatum of older subjects (Old_Str). This module is upregulated in the elderly group but significantly downregulated in both Alzheimer’s disease (AD) and Parkinson’s disease (PD) groups. These findings highlight distinct glycosylation patterns and offer insights into the potential molecular mechanisms driving neurodegenerative diseases and aging processes.

### N-GlycoMiner: A Comprehensive Database of Experimentally and Theoretically Characterized N-Glycoproteomes

There is a significant shortage of comprehensive intact N-glycopeptide databases for examining protein glycosylation level. Existing resources, such as N-GlycositeAtlas and GPnotebook, have laid foundational groundwork, but they do not cover the full scope of N-glycoproteomic diversity across species and disease contexts^40, 41^. To address this gap, we developed N-GlycoMiner, a database (accessible at www.NGlycoMiner.com) that consolidates site-specific glycopeptides from experimental mouse N-glycoproteome data in this study, as well as from 60 key publications spanning the last decade (2014–2024) (Figure 6 and Supplementary Data 7).

**Figure 6.**
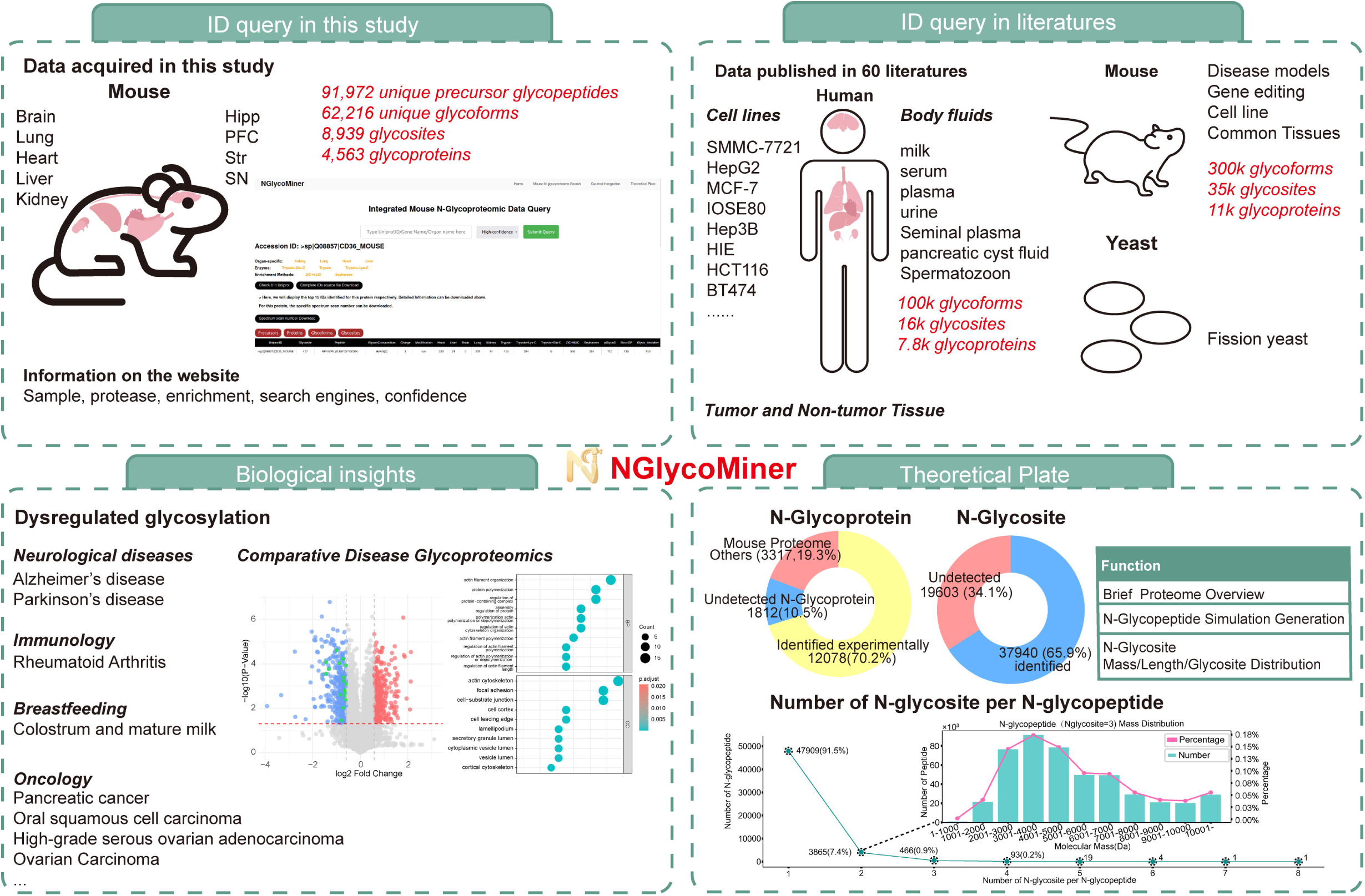
An integrative platform for N-Glycoproteomics ID query.

For the data in this study, the website showed detailed information for each N-glycoprotein, including tissue specific expression, proteases used, enrichment methods, spectrum scan number, multi-leveled identification results, search engines and identification confidence levels. The protein structures predicted by AlphaFold2 and the identified glycosites and glycan compositions were visualized. A total of 4,563 mouse glycoproteins in this study with tissue-specific expression profiles were showed. The spatial-temporal and disease-related expression data in this study can be queried.

Our study adds to the existing body of knowledge with 91,972 precursor glycopeptides, 62,216 unique glycoforms, 8,939 glycosites, and 4,563 glycoproteins identified in mouse samples from various organs, including the brain, heart, liver, lung, and kidney. Combined with literature data, our platform offers a total of 310,416 glycoforms, 12,076 glycoproteins, and 37,931 glycosites for mouse samples, and 107,459 glycoforms, 8,007 glycoproteins, and 16,624 glycosites for human samples, making it the most comprehensive N-glycoproteome resource to date. Notably, this study contributed an additional 40,216 glycoforms, 2,213 unique glycosites, and 288 glycoproteins beyond previously reported literature findings (Figure S21).

In addition to facilitating glycoproteome searches, N-GlycoMiner offers insights into disease-associated dysregulated glycosylation patterns. Neurological diseases, such as Alzheimer’s and Parkinson’s disease, are represented alongside cancers and immune disorders, highlighting the role of N-glycoproteins as potential biomarkers. Comparative analysis of disease-specific glycopeptides reveals functional annotations and pathway associations, offering valuable data for further biomarker validation and therapeutic target exploration^42-44^

Moreover, our database includes theoretical analyses to predict possible N-glycosylation sites and N-glycoproteins using the canonical sequon N-X-S/T/C (where X ≠ P) (Figure S22). For example, in silico analysis of the mouse proteome predicts 57,543 N-glycosites and 13,890 N-glycoproteins. Based on our database, about 66% of total putative N-glycosites and 70.2% of total proteins were supported by experimental evidence. Additionally, around 92% of experimentally detected tryptic glycopeptides feature a single glycosylation site, simplifying site localization for N-glycosylation. For complex glycopeptides or those with multiple glycosylation sites, alternative enzymes, such as Glu-C, are recommended on the website to aid in detection (Figure S22g). In summary, N-GlycoMiner consolidates the largest dataset for mouse and human N-glycoproteomes, bridging experimental data with theoretical predictions.

## Discussion

Despite many advances in glycoproteomics technology and search engines^45, 46^, the identification depth and accuracy of N-glycoproteomics are still insufficient. In this study, although we provided a comprehensive, high-quality glycoproteomics dataset and performed joint retrieval with four widely used glycoproteomics search engines, the interpretation rate of the glycopeptide spectra was as low as only 11.6%. Moreover, among the 1,041,225 GPSMs identified, only 160,928 were consistently identified by all the four engines, while 46.8% of GPSMs had low confidence, highlighting the limitations of current glycoproteomics technology.

The study found that over 88.4% of GPSMs remained unidentified. Many spectra had low-quality spectra with minimal ion information, likely due to insufficient fragmentation efficiency and low glycoform abundance, making identification challenging. Other spectra, despite exhibiting high-quality fragmentation patterns, were still not identified, highlighting limitations in current algorithms (Figure S11). Several factors may have influence on the low identification rate. Specific glycan modifications (e.g., phosphorylation, sulfation, O-acetylation) can alter spectral profiles, complicating software-based identification^47^. Rare or complex glycans, such as polysialic acid and poly-LacNAc, may be absent from glycan databases, leading to missed identifications^48, 49^. Mixed spectra, where fragment ions come from multiple glycopeptides, can also confuse software^19^. Additionally, the presence of O-glycopeptides, which were not specifically removed, may contribute to unidentified spectra. To improve spectra quality, optimizing fragmentation conditions—such as adjusting HCD energy or using techniques like AI-ETD, EThcD, or ion mobility— could enhance spectral resolution. AI-assisted spectral library prediction, like DeepGP, has also shown promise in improving identification accuracy^32, 33^. These advancements in MS and AI-driven analysis offer the potential to significantly enhance glycopeptide identification in future studies.

Artificial intelligence is a promising means to enhance the ability of search algorithms^31, 32, 33^.The large-scale, high-quality and graded glycopeptide datasets generated in this study have been shown to enhance AI-driven glycopeptide spectral prediction, retention time estimation, and glycoproteomics software development. We believe our data will continue advancing glycoproteomics tools, as seen in our 2017 study.

We reported a largest atlas of moue N-glycoproteome consisting of 62,216 unique glycoforms, 8,939 glycosites, and 4,563 glycoproteins. Compared to the glycoproteins in UniProt, this study additionally identified 2,847 glycoproteins and 5,177 glycosites, significantly expanding the mouse N-glycoproteome database. The number of GPSMs and glycoforms in this study increased by 13-fold and 6.2-fold, respectively, compared to 2017 (pGlyco2)^19^.

Analysis of this dataset revealed complex glycosylation microheterogeneity, particularly tissue-specific variations. The brain displayed higher microheterogeneity compared to other organs, reflecting its unique physiological requirements. Comparative analysis of the glycoprotein functions across tissues showed that brain-specific glycoproteins are primarily involved in neurodevelopment and synaptic function, while liver, heart, kidney, and lung glycoproteins are associated with metabolism, homeostasis, and cellular functions unique to each organ.

We established a comprehensive database for mouse and human N-glycoproteome, N-GlycoMiner, which addresses the significant gap in site-specific N-glycoproteomics databases. This resource in total encompasses 310,416 glycoforms, 12,076 glycoproteins, and 37,931 glycosites for mouse, as well as 107,459 glycoforms, 8,007 glycoproteins, and 16,624 glycosites for human. Notably, this dataset represents 66% of theoretical prediction sites for mice. These extensive datasets contribute to a thorough understanding of the biological relevance of glycosylation. Additionally, N-GlycoMiner highlights disease-associated dysregulated glycosylation patterns, particularly in neurological diseases and cancers, providing valuable information for biomarker validation.

In conclusion, our work delivers a robust and valuable resource for glycosylation research. The large-scale, ultra-deep dataset not only enhances our understanding of glycosylation microheterogeneity and tissue specificity but also provides critical insights into how glycosylation is dysregulated in aging and disease contexts, especially in neurodegeneration. This resource lays the groundwork for future studies that will further explore glycosylation’s functional roles and therapeutic potential in health and disease.

## Methods

### Mouse

C57BL/6 mice were housed in individually ventilated cages with free access to water and standard mouse chow, following ethical regulations approved by the Ethics Committee of Fudan University, China. At 8 weeks of age, both male and female mice were euthanized using CO2. The animals were then perfused with 50 ml of precooled phosphate-buffered saline. Following perfusion, tissues including the brain, lung, kidney, liver and heart were dissected, snap-frozen in liquid nitrogen, and stored at −80 °C. For Alzheimer’s disease (9 months old) and Parkinson’s disease (3 months old) studies, mice were sourced from Zhishan Institute of Health Medicine Co., Ltd. (Beijing, China). Brain regions including the hippocampus (Hipp), medial prefrontal cortex (mPFC), substantia nigra (SN), and striatum (STR) were isolated for analysis. Both male and female C57BL/6 mice were used in this study. Sex was not considered as a variable in the study design or analysis. All animal procedures were approved by the Ethics Committee of Fudan University, China.

### Protein extraction

Tissues were homogenized in lysis buffer (4% SDS, 0.1 M Tris/HCl, pH 7.6) containing a protease inhibitor cocktail using a high-throughput tissue grinder (ONEBIO, Shanghai, China) at 65 Hz for 60 seconds. The homogenates were then sonicated and clarified by centrifugation at 14,000 rpm for 40 minutes. Protein concentrations were measured using the BCA assay (Pierce, Rockford, IL).

### Protein digestion

Proteins were first reduced with 10 mM DTT at 37°C for 60 minutes, followed by alkylation in the dark with 20 mM iodoacetamide (IAA) at room temperature for 30 minutes. After carbamidomethylation, six volumes of acetone were added to precipitate the proteins at −20°C for at least 3 hours. The precipitates were then completely dissolved in a denaturing buffer (8 M urea in 50 mM NH4HCO3) and diluted 10-fold with 50 mM NH4HCO3. For digestion, trypsin, trypsin coupled with Lys-C, or trypsin coupled with Glu-C were added at a ratio of 50:1 (protein to enzyme) and incubated for 12-16 hours. The reactions were halted by heating at 95°C for 10 minutes. The digested samples were then acidified and centrifuged at 14,000 × g for 10 minutes. The supernatants were desalted using Sep-Pak C18 columns (Waters Corporation, Milford, MA, USA) according to the manufacturer’s instructions, then dried by vacuum centrifugation, and stored at −80°C for further use

### Glycopeptide enrichment

Two enrichment methods, ZIC-HILIC and Sepharose CL-4B, were employed in this study. For ZIC-HILIC enrichment, digested peptides (1 mg) were dissolved in a loading buffer (80% (v/v) acetonitrile and 1% (v/v) trifluoroacetic acid) and loaded onto a micro-column containing 30 mg of ZIC-HILIC medium (5 μm, Merck Millipore, Darmstadt, Germany). After washing three times with the loading buffer, the retained analytes were eluted with 1 mL of 0.1% trifluoroacetic acid, followed by 20 μL of 25 mM ammonium bicarbonate and 25 μL of 50% acetonitrile.

For glycopeptide enrichment using Sepharose CL-4B, digested peptides (1 mg) were mixed with 100 μL of Sepharose CL-4B in 500 μL of an organic solvent mixture consisting of butanol/ethanol/water (4:1:1 by volume). The mixture was gently shaken for 60 minutes, after which the resins were washed three times with the organic solvent. The resins were then incubated with an aqueous solvent of ethanol/water (1:1 by volume) for 30 minutes, and the solution phase was recovered. The enriched glycopeptides were subsequently dried by vacuum centrifugation and stored at −80°C for further use

Liquid chromatography-mass spectrometry (LC-MS). Glycopeptides extracted from mouse tissues were analyzed using an Orbitrap Fusion mass spectrometer equipped with a Proxeon EASY-nLC II liquid chromatography pump (Thermo Fisher Scientific). The analysis utilized a reversed-phase analytical column (Thermo Scientific, C18, 500 mm × 75 µm, 3 µm). Mobile phase A consisted of water with 0.2% (v/v) formic acid, while mobile phase B was acetonitrile with 0.2% (v/v) formic acid. Samples were loaded onto the column at a flow rate of 2 μL/min for 3 minutes. The flow rate during the run was maintained at 200 nL/min with the following linear gradient: 1% to 40% B over 345 minutes, 40% to 90% B over 3 minutes, followed by a 5-minute re-equilibration at 1% B. Internal mass calibration was performed using a defined lock mass, m/z 445.12003. Survey scans of peptide precursors in the 350 to 2000 m/z range were conducted at 120K resolution, with a target ion count of 5 × 10^5 and a maximum injection time of 50 ms, using the Orbitrap detector. The microscan setting was 1, and the s-lens RF level was set to 60. Tandem MS (MS2) was performed with an isolation window of 4 Th using the quadrupole, HCD fragmentation with stepped collision energies of 20%, 30%, and 40%, and MS2 analysis in the Orbitrap at a resolution of 15,000. The MS2 ion count target was set to 5 × 10^5 with a maximum injection time of 250 ms. Only precursors with charge states of 2 to 6 were selected for MS2. Dynamic exclusion was set to 30 seconds with a 10-ppm tolerance around the selected precursor and its isotopes. Monoisotopic precursor selection was enabled. The instrument operated in top-speed mode with 3-second cycles, continuously performing MS2 events until either the list of non-excluded precursors was exhausted or 5 seconds had elapsed, whichever occurred first.

Glycopeptides from Alzheimer’s disease and Parkinson’s disease samples were analyzed using an Orbitrap Exploris 480 mass spectrometer equipped with a FAIMSpro (High Field Asymmetric-Waveform Ion-Mobility Spectrometry) interface, coupled with an Easy-nLC 1200 system (all from Thermo Fisher, San Jose, CA). Separation was achieved using 40-cm high-performance liquid chromatography (HPLC) columns (75 μm inner diameter monolith column, Kyoto Monotech, Kyoto, Japan). For each LC-MS/MS analysis, approximately 1000 ng of peptides were injected for a 120-minute gradient. Peptides were loaded in buffer A (0.1% formic acid) and separated with a linear gradient at a flow rate of 300 nL/min: from 5% to 20% buffer B over 90 minutes, from 20% to 55% B over 20 minutes, followed by a 10-minute re-equilibration at 5% buffer B.

MS spectra were acquired using a cycle time mode with multiple compensation voltages (CVs) applied. The CVs used were -45V, -50V, and -65V, with a cycle time of 1 second for each CV. The normalized AGC (Automatic Gain Control) target for full scan MS spectra was set to 300%, with an injection time of 50 ms. The m/z range was set from 350 to 2000, and the resolution was 120,000 at m/z 200. The RF lens was set to 50%. Fragmentation of precursor ions was performed using higher-energy collisional dissociation (HCD) with normalized collision energies of 20%, 30%, and 40%. MS/MS scans were conducted at a resolution of 15,000 at m/z 200, with an AGC target of 50% and a maximum injection time of 100 ms.

Database searches for N-glycoproteomics. Mouse protein databases containing reviewed and canonical sequences were downloaded from UniProt. The raw data were processed using various software, all with a precursor tolerance of 10 ppm for MS1 and 20 ppm for MS2. Specific cleavage sites were defined for trypsin and trypsin coupled with Lys-C at lysine (K) and arginine (R), and for trypsin coupled with Glu-C at lysine (K), arginine (R), aspartic acid (D), and glutamic acid (E), allowing for up to two missed cleavages. Carbamidomethylation of cysteine (Carbamidomethyl[C]) was set as a fixed modification, while oxidation of methionine (Oxidation[M]) and acetylation of the protein N-terminus (Acetyl [ProteinN-term]) were set as variable modifications. The fragmentation type was set to HCD. The false discovery rate (FDR) was controlled at 1% for each search engine.

1. pGlyco3^20^: The pGlyco version used was pGlyco3.0.rc2, downloaded from [http://pfind.org/software/pGlyco/index.html#Downloads]. Fragmentation of HCD were selected. A maximum of two variable modifications on peptides was allowed. The peptide length was set between 6 and 40 amino acids, with a mass range from 600 to 4000 Da. The glycan database selected was pGlyco-N-Mouse.gdb, with no additional glycan modifications applied. The setting for pGlycosite was Localized Glycan Must be in GDB. All other parameters were left at their default settings. The extracted ion chromatography (XIC) areas for monoisotopic glycopeptide precursors as reported by pGlyco3 were used for quantification.
2. StrucGP^27^: Version 1.2.0 of StrucGP was employed for this study. A total of 17 branch structures from the default branch structure database were selected. Collision energies were set at 20, 30, and 40 in this study while both low and high energy settings at 30 were used for database search as recommended by the software developer. The top 300 fragment ions were considered, and at least two oxonium ions (138.055 and 204.0866) from the top 20 fragment ions in the MS/MS spectra were used to extract the MS/MS spectra for intact glycopeptides. A minimum of five peptide fragments was required. Identification results were filtered to a 1% false discovery rate (FDR) for both peptide sequences and glycans, using the decoy peptide database and decoy spectra methods for estimation, respectively. All other parameters were set to default.
3. MSFragger-Glyco^26^: The FragPipe version 19.2-build28, MSFragger version 3.7, IonQuant version 1.9.4, and Philosopher version 5.0.0-RC14 were used. A workflow of glyco-N-HCD were loaded. Glycan database of Mouse_N-glycans-1670-pGlyco was selected. All other parameters were left at their default settings. Glyco-N-LFQ workflow were used for quantification.
4. Glyco-Decipher^28^: Version 1.0.4 of Glyco-Decipher was used. Spectrum expansion was enabled, and the GlyTouCan database was utilized for glycan matching. The monosaccharide stepping method was employed to determine the composition of modified glycans that did not match any entries in the database. All other parameters were set to default. Label free quantification without match between run were used.

### Data Analysis

All data processing and visualization were conducted using Python (version 3.7) and R (version 4.2.2). Quantitative analysis of tissue-specific N-glycopeptides in mice was performed using extracted ion chromatography (XIC) areas for monoisotopic glycopeptide precursors, as reported by pGlyco3 (Supplementary Data 4). For precursors with differing XIC profiles due to isomer variations, the maximum values were utilized. For each raw file, the XIC areas of all precursor glycopeptides corresponding to the same unique glycoform were summed for quantification. The summed areas of all unique glycoforms, corresponding to the same unique glycosites, glycan compositions, or glycoproteins, were calculated. Finally, total intensity normalization methods were applied to ensure comparability across different experimental runs. The normalized table were combined to establish the whole expression matrix for all samples. Missing values were imputed using 0 followed by log2 transformation for further analysis.

Distribution information of the N-glycosite on protein secondary structure were obtained from Alphafold2 predictions and DSSP (Dictionary of Protein Secondary Structure) annotations^50^. PCA was implemented with the prcomp function in R script and the first two principal components were visualized using the factoextra package. Unsupervised hierarchical clustering was performed on the expression matrix following z-score. Pearson correlation analysis was performed based on normalized glycoprotein expression between each protein groups co-detected in our glycoproteomics data. Putative co-expression pairs were retained if their PCCs were above 0.6. Known protein-protein interaction (PPI) were retrieved from STRING database (https://stringdb.org/api) and visualized by Cytoscape (version 3.8.0). GO and KEGG pathway enrichment analyses were performed using g:Profiler^51^. Pathways with p-value threshold of <0.05 were considered significantly enriched.

Mouse brain region-specific quantification data was obtained from XIC areas of monoisotopic glycopeptide precursors, as reported by pGlyco3 (Supplementary Data 6). The rows with only one non-zero value out of three replicates are removed. The combined expression matrix undergoes KNN imputation with n_neighbors=2 to fill in the missing values. After this, the data was further processed by performing total intensity normalization followed by log2 transformation for further analysis. Unpaired Students *t*-test with two-tailed were used for the significantly changed glycoproteome analysis. Volcano plot with fold change (log2 ratio)>1 and *p* value <0.05 were used.

### WGCNA

To gain system-level insights into brain N-glycoproteome changes in neurodegeneration and aging models, we conducted WGCNA to construct a co-regulation network for N-glycosites. Initially, we preprocessed the quantification data followed by hierarchical clustering to identify outliers. We selected a soft-threshold power of 9 and minModuleSize of 10 to ensure a scale-free topology. The resulting modules were then correlated with binary traits relevant to neurodegeneration and aging, revealing key modules significantly associated with these traits. “Hub glycosites” were defined as those with intramodular connectivity (k within values) within top 10% of the respective module. For glycan composition matrix, a consistent approach was applied with a soft threshold set to 7.

### Website construction

A user-friendly interface was developed to browse through the database and generate theoretical N-glycopeptides for research, available at www.NGlycoMiner.com. The site is built on the Django web framework, utilizing MySQL as the storage database, written in the Python scripting language (v 3.8) for the backend. The frontend features a responsive design crafted with HTML, CSS, and JavaScript. The application is deployed with Nginx as a reverse proxy and uWSGI. A step-by-step PowerPoint presentation to guide users in navigating the website were provided in Supplementary Data 8 and on website.

### The prediction of MS/MS spectra and retention time using AI models

The datasets utilized for training and testing were derived from subsets of pGlyco3, stratified based on different confidence levels. The MGF files were also generated by pGlyco3. The prediction of MS/MS spectra was conducted using DeepGP and DeepGlyco model using default parameters ^32, 33^. Glycopeptides retention time prediction was executed by the DeepGP model.

### Western blot analysis of glycosylation modifications on CD36

Male C57BL/6 mice were anesthetized using Avertin. Heart, liver, brain, lung, and kidney tissues were quickly dissected and snap-frozen in liquid nitrogen for subsequent analysis. Total proteins were extracted from these tissues using an ice-cold mild lysis buffer (Sigma, C2978) supplemented with protease inhibitors. Protein concentrations were determined using a BCA protein assay kit (Thermo Fisher Scientific, 23227). Native proteins were heated under the denatured conditions. Proteins were separated by SDS-PAGE and transferred onto PVDF membranes (Millipore, IPVH00010). The membranes were incubated overnight at 4°C with primary antibodies against CD36 (Abcam, ab252922) at a 1:1000 dilution and GAPDH (Proteintech, 10494-1-AP) at a 1:5000 dilution. Following washing with TBST, the membranes were incubated with HRP-conjugated secondary antibody (Anti-Rabbit IgG-HRP, Beyotime, A0208) at a 1:5000 dilution for 1 hour at room temperature. Protein bands were detected using an ECL reagent (Epizyme, SQ202) and visualized with the ChemiDoc XRS+ imaging system (Bio-Rad). For deglycosylated treatment, 50 μg of tissue proteins were treated with 0.2 μL of PNGase F (New England Biolabs, P0704S) at 37°C, 800 rpm for 18 hours to remove N-glycans. Following treatment, samples were analysed by Western Blot to detect both glycosylated and non-glycosylated CD36 expression.

### Statistics and Reproducibility

For analyses involving mouse tissues (brain, lung, kidney, liver, and heart), five LC-MS/MS replicates were used. Mice were randomly assigned to experimental groups prior to Alzheimer’s disease (AD) and Parkinson’s disease (PD) modeling. No statistical method was used to predetermine sample size. For the AD (9 months old) and PD (3 months old) brain region studies, three biological replicates were included per group. Statistical significance for group comparisons was assessed using two-tailed unpaired Student’s t-tests, as detailed in the figure legends and in the website. All key experiments were independently repeated with consistent results. The statistical tests used are indicated in the figure legends.

## Data availability

All raw mass spectrometry data have been deposited in the iProX partner repository under the dataset identifier IPX0011099000^52^. The data are publicly accessible. Source data are provided with this paper.

## Acknowledgements

This work was supported by several funding sources, including the National Key Research and Development Program (2021YFA1301601, 2022YFC2503904), the Science and Technology Innovation 2030 Major Projects (2022ZD0211600), the National Natural Science Foundation of China (82272174, 81827901, 32101026, 32471504, 22201217, 62475179, 32200778, 32471014), the Science and Technology Innovation Action Plan of STCSM (22S31901900) and the innovative research team of high-level local university in Shanghai, NHC Key Laboratory of Glycoconjugates Research, as well as the Brain Health Youth Fund - Precision Diagnosis and Treatment of Alzheimer’s Disease Research in 2024. Natural Science Foundation of Jiangsu Province (NO. BK20220494), Suzhou International Joint Laboratory for Diagnosis and Treatment of Brain Diseases, the Priority Academic Program Development of Jiangsu Higher Education Institutes (PAPD), the Project of MOE Key Laboratory of Geriatric Diseases and Immunology (No. KJS2501). The Clinical Research Center of Neurological Disease in The Second Affiliated Hospital of Soochow University (NO. ND2022A04). We also acknowledge the proteomics and mass spectrometry platform at the Institutes of Biomedical Sciences, Fudan University. This paper is dedicated to the memory of Professor Pengyuan Yang (1949–2021), who passed away during the revision of the work.

## Author contributions

H.L.S. and P.F. supervised the project. P.F. conducted the experiments and data analysis. X.M.Y conducted the data analysis. H.Y.J., Q.S., Y.N.L., W.W.Z., W.M.Z. and Z.X.L. contributed to data analysis. M.Y.D. and Q.F.C. contributed to the wet experiments. W.J.K. and P.Y.Y. contributed to the manuscript. P. F. and H.L.S. wrote the manuscript with help from all other authors.

