## Supplementary information for "Ultradeep N-glycoproteome Atlas of Mouse Reveals Spatiotemporal Signatures of Brain Aging and Neurodegenerative Diseases"

Correspondence：

Supplementary Note 1. Evaluation of Enzymatic Digestion and Enrichment Methods in Large-Scale Glycoproteomics

Supplementary Note 2. Comparative Analysis of N-Glycoproteomics Software Tools

Supplementary Note 3. Large-Scale and High-Quality Glycoproteomic Spectra Enhance Prediction Performance in Existing AI Models for N-Glycopeptide Tandem Spectra and Retention Time.

Figure S1. Overview of Collected Scans and Identification Annotation Process.

Figure S2. Quality Control Metrics for N-Glycoproteomic Analysis.

Figure S3: Comparison of Glycopeptide Analysis Using Different Proteases.

Figure S4. Comparison of Glycopeptide and Glycan Compositions Between Sepharose and ZIC-HILIC Enrichment Methods.

Figure S5. Saturation Curves of Unique Glycoform Identification Across Various Tissues.

Figure S6: Comprehensive Analysis of Glycoproteomic Data Across Different Software Tools.

Figure S7. Comparison of Glycoproteomics Analysis Results Across Different Software Tools.

Figure S8. Comparison of Quantitative Glycoproteomics Data Between Brain and Heart Samples Using Different Software with Trypsin and Trypsin+Lys-C Digestion.

Figure S9. Analysis of Inconsistent Precursor Identifications Across Four Software Tools.

Figure S10. Enhancement of AI Training Performance with Datasets of Varying Quality.

Figure S11. Example Spectra for Unidentified Spectra.

Figure S12. Comparison and Identification of Glycoforms and Glycoproteins Across Mouse Tissues and Enzyme Treatments.

Figure S13: Comparison of Glycoproteomic Metrics(pGlyco3 Results) Across Different Tissues and Enrichment Methods.

Figure S14: Analysis of Glycoproteomic Data Across Multiple Tissues and Structural Insights.

Figure S15. Western Blot Analysis and Glycoform Identification of CD36 Glycoprotein Across Different Tissues.

Figure S16: Comparative Analysis of Glycosylation Patterns Across Different Tissues.

Figure S17. Volcano Plots Showing Differential Protein Expression in Various Brain Regions of Alzheimer's Disease (AD) and Parkinson's Disease (PD) Patients.

Figure S18. Volcano Plots Depicting Differential Protein Expression Associated with Aging in Different Brain Regions.

Figure S19. Module-Trait Relationships for Glycan Composition Analysis.

Figure S20. Hierarchical Clustering and Functional Enrichment Analysis of Glycosite Modules.

Figure S21. Venn Diagrams Illustrating the Overlap Between Curated Literature and Experimental Mouse Data.

Figure S22. Analysis of N-Glycopeptide Mass Distribution and Glycosite Reduction Across Different Enzyme Digestions.

**Supplementary Note 1.** **Evaluation of enzymatic digestion and enrichment methods in large-scale glycoproteomics**

We systematically evaluated the performance of three enzymatic digestion strategies and two glycopeptide enrichment methods and integrated them to enhance glycoproteome coverage and improve the reliability of large-scale glycosylation analysis. Our findings provide a detailed assessment of their complementarity and biases, offering valuable insights into optimizing glycoproteomics workflows.

**1. Enzymatic digestion strategies and their complementarity**

**(1) Complementarity of trypsin and Trypsin+Lys-C**: In our large-scale mouse dataset, identification numbers for each enzyme plateaued after multiple runs, indicating a saturation limit (Figure S3a). While both enzymes cleave at lysine (K) and arginine (R), their complementarity is evident: trypsin uniquely identified 8503 glycopeptides, while Trypsin+Lys-C identified 5526 (Figure S3b). Trypsin alone yielded slightly more glycopeptides than the trypsin + Lys-C combination but had a higher missed cleavage ratio (Figure S3c). Both enzymes showed a similar glycosylation site distribution, with a preference near the N-terminal (Figure S3d).

**(2) Contribution of Trypsin + Glu-C**: This combination provided an additional 6151 unique identifications compared to trypsin alone, particularly enriching peptides ending with glutamic acid (E) or aspartic acid (D) (Figure S3b). Although its total identifications were slightly lower, likely due to shorter peptides generated by Glu-C, it contributed valuable complementary coverage.

**(3) Peptide molecular weight distribution**: Trypsin and trypsin + Lys-C produced nearly identical molecular weight distributions, while trypsin + Glu-C yielded shorter peptides due to Glu-C’s cleavage specificity (Figure S3e, f), supporting the robustness of our approach.

**(4) Necessity of multi-enzyme digestion**: In our previous published work^1^, we systematically compared the performance of multiple proteases for glycosite identification by removing glycans with PNGase F and identifying deglycosylated peptides based on the 0.98 Da mass shift. This study demonstrated that the combination of three enzymes (trypsin, Lys-C, and Glu-C) provided the highest glycosite coverage. Other proteases, such as Chymotrypsin, Glu-C, and Pepsin, yielded significantly fewer identifications. Based on these results, we chose these three enzymes, and the current study further validates their complementarity and reliability for comprehensive and large-scale glycoproteomics analysis.

**2. Comparative analysis of glycopeptide enrichment methods**

**(1) Differences in enrichment principles**: ZIC-HILIC employs a mixed-mode mechanism for separating and enriching glycopeptides, involving hydrophilic partitioning, electrostatic interactions, dipole interactions, adsorption, and hydrogen bonding. This mixed-mode mechanism provides broader coverage for glycopeptide enrichment, as demonstrated in previous studies^2^. Sepharose CL-4B primarily operates based on hydrophilic interaction chromatography (HILIC), where glycopeptides are enriched through hydrogen bonding with polar groups in glycopeptides. These distinct mechanisms contribute to differences in their enrichment performance.

**(2) Glycopeptide identification**: In our ultra-deep dataset, both enrichment methods achieved saturation in glycopeptide identification (Figure S4a), enabling a comprehensive comparison of the differences between the two methods. This saturation demonstrates that while both methods are effective, they provide complementary benefits for glycopeptide enrichment. ZIC-HILIC yielded higher glycopeptide identification overall but showed significant complementarity with Sepharose CL-4B (Figure S4b). This complementarity underscores the importance of combining multiple enrichment methods to capture the full range of glycopeptides present in the sample. Using only one method cannot fully enrich all glycopeptides, highlighting the necessity of complementary approaches.

**(3) Enrichment efficiency**: The enrichment efficiency of ZIC-HILIC was consistently high (greater than 70%), as shown in Figure S4c, while Sepharose CL-4B exhibited more variable efficiency, ranging from 30% to 80%. This variability suggests that while ZIC-HILIC is more reliable for large-scale analyses, Sepharose CL-4B may still capture certain glycopeptides that are not efficiently enriched by ZIC-HILIC.

**(4) Glycan identification**: ZIC-HILIC covered a wider range of glycan compositions, whereas Sepharose CL-4B contributed only a small number of unique identifications (**Figure S4b**). Further analysis of glycan types revealed that Sepharose CL-4B showed a slightly higher proportion of high-mannose and paucimannose glycans (Figure S4e). For sialylated glycans, the two methods were comparable, but ZIC-HILIC enriched a higher proportion of fucosylated glycans than Sepharose CL-4B.

Our study demonstrates that both ZIC-HILIC and Sepharose CL-4B have distinct advantages and complement each other in glycopeptide enrichment. The combined use of these methods enables more comprehensive glycopeptide coverage, enhancing the depth and reliability of glycosylation analysis.

**Supplementary Note 2. Comparative analysis of N-glycoproteomics software tools**

In this study, we conducted a comprehensive comparison of four widely used glycoproteomics software tools—pGlyco3, MSFragger-Glyco, Glyco-decipher, and StrucGP—to assess their biases and performance across different identification and quantification levels. Our findings highlight critical trade-offs among these tools, offering insights into their respective strengths and limitations.

**(1) Bias in identification across different levels.** At the GPSM, precursor, and glycoform levels (Figure 2 and Figure S6), the rankings of identification numbers are almost consistent across all tools, with Glyco-decipher achieving the highest, followed by MSFragger-Glyco, pGlyco3, and StrucGP, which has the lowest. However, at other identification levels, including glycosites, glycoproteins and glycan compositions, the results vary significantly. pGlyco3 exhibits the lowest identification numbers for glycosites and glycoproteins but performs moderately well in other categories, notably surpassing MSFragger-Glyco in glycan identification. StrucGP, on the other hand, performs comparably to MSFragger-Glyco and Glyco-decipher at the glycosite and glycoprotein levels. This analysis underscores the trade-offs between the tools in their ability to identify glycoproteins, glycosites, and glycans, highlighting the influence of their design and analytical focus on identification performance.

**(2) Identification consistency analysis.** For the spectra identified by all four tools (191,981 in Figure 2b), 160,928 spectra were consistently identified as the same precursors by all tools, while 31,053 spectra showed inconsistencies. MSFragger-Glyco contributed to the highest proportion of inconsistent identifications (35.6%), followed by Glyco-decipher (18.1%) and StrucGP (12.8%). In contrast, pGlyco3 had the lowest inconsistency rate (4.2%) (Figure S9a). For the spectra identified by any three of the four tools, MSFragger-Glyco still exhibited the largest inconsistency (13.3%), followed by Glyco-decipher (3.7%) and pGlyco3 (1.7%) (Figure S9b). These results suggest that pGlyco3 is the most reliable tool considering consistent spectrum identification. On the other hand, MSFragger-Glyco identifies a larger number of spectra with greater inconsistency. The high inconsistency rate of MSFragger-Glyco indicates that while it has higher sensitivity, it may also lead to more false positives. These differences highlight the varying strengths and biases of these tools, with pGlyco3 excelling in reliability and MSFragger-Glyco providing a broader range of identifications at the expense of consistency.

**(3) Bias in peptide and glycan features.** We specifically analyzed the distribution of peptide length (number of amino acids), peptide molecular weight, glycan length (number of monosaccharides) and glycan types across the four software tools (Figure S7). StructGP consistently identifies shorter peptides and glycans, whereas MSFragger-Glyco shows a preference for longer peptides, and Glyco-decipher tends to identify longer glycans. Notably, pGlyco3’s glycan-length bias was influenced by the exclusion of glycan structures above 20 monosaccharides, as we used the pGlyco-N-Mouse database instead of pGlyco-N-Mouse-Large. In terms of glycan types, StructGP is biased toward high-mannose and pauci-mannose glycans, with the lowest identification of sialylated glycans, indicating its limited ability to detect complex glycan structures. On the other hand, MSFragger-Glyco exhibits the highest sialic acid content in its identifications. Meanwhile, both pGlyco3 and Glyco-decipher demonstrate a stronger focus on fucosylated glycan identifications. Collectively, these observations highlight distinct differences in the analytical focus and limitations of the tools, emphasizing the need to carefully select software based on the specific requirements of a glycoproteomics study.

**(4) Quantification Bias: Heart vs. Brain Samples.** For quantification evaluation, we compared pGlyco3, Glyco-decipher, and MSFragger-Glyco, as these are the only tools supporting quantitative analysis. We found that the differentially expressed glycoproteins and glycosites showed a high overlap in up-regulated and down-regulated molecules, indicating a high consistency in quantification between the three software tools at these levels (Figure S8a). However, at the glycan and site-specific glycoform levels, the overlap in quantification results was much lower compared to glycoproteins and glycosites, indicating lower consistency in quantification at these levels.

We further conducted a correlation analysis of the quantification values among the three software tools using their commonly identified and quantified glycopeptides (Figure S8b). The highly positive correlations in the quantification of glycoproteins and glycosites (median PCC above 0.78) were revealed, suggesting that the three software platforms agree well in quantifying these levels of data. However, the correlation analysis for glycans showed more variability, with lower median PCC (median PCC ranges 0.52-0.67). This suggests that the software tools are generally consistent in quantifying glycoproteins and glycosites, but less agreement at the glycans levels. This inconsistency could be due to differences in how each tool processes and interprets glycan-related data. This lower consistency in glycans’ and glycoforms’ quantification may lead to divergent interpretations in subsequent studies of glycosylation's biological functions.

Overall, these findings emphasize the need for caution when comparing glycan and glycoform data across different software platforms, particularly in studies focused on the functional implications of glycosylation. Further improvements in glycan quantification algorithms and validation with non-omics techniques could help address these discrepancies.

**（5）Comparison of the four software tools from their identification strategies, glycan and peptide scoring and false positive control.** Each software tool utilizes a distinct search strategy for glycoproteomic identification, which impacts their performance in glycan and peptide identification and false positive control.

pGlyco3 adopts a glycan-first search strategy, which prioritizes the identification of N-glycopeptides by first identifying glycans using ion-indexing techniques, followed by peptide identification^3, 4^. This approach is integrated with scoring systems for glycans, peptides, and glycopeptides. A notable feature of pGlyco3 is its rigorous false discovery rate (FDR) control, which applies across all levels—glycans, peptides, and glycopeptides. pGlyco3 used glycan database combining N-glycans from Glycome-DB, as well as the three largest substructures of high-mannose, hybrid, and complex N-glycans. This glycan-first approach and FDR control allow pGlyco3 to outperform the other tools in terms of accuracy though the strict glycan-level quality control may reduce sensitivity (Figure 2, S6 and S9).

StrucGP uses a modular approach, dividing N-glycans into three modules, with each identified based on distinct Y ion patterns or combinations of B/Y ions^5^. This methodology facilitates detailed glycan structure determination. FDR estimations for both the peptide and glycan portions of glycopeptides are performed using both decoy database and decoy spectrum approaches. While StrucGP performs fewer glycan identifications compared to other tools, it offers similar performance to MSFragger-Glyco and Glyco-Decipher in terms of glycosite and glycoprotein identification (Figure S6). This is likely because StrucGP requires more ion information for detailed glycan structure analysis, which makes it more prone to identifying shorter peptides and glycans, especially high-mannose glycan types (Figure S7).

MSFragger utilizes an open mass offset search strategy, where glycopeptide spectra are analyzed based on possible glycan compositions derived from mass offsets. It performs deisotoping and decharging of spectra prior to analysis, improving the sensitivity of the tool, particularly for high-mass glycopeptides (Figure S7). However, MSFragger does not place a strong emphasis on glycan-level quality control, which may lead to less precise glycan identification compared to other tools^3^. The high sensitivity of MSFragger often results in a larger number of identifications, but it has the lowest consistency in identifications compared to the other software (Figure 2, S6 and S9).

Glyco-Decipher employs a glycan database-independent peptide matching strategy, which uses shared fragmentation patterns of peptide backbones in glycopeptides to improve spectrum interpretation^6^. This technique enables Glyco-Decipher to extend its glycopeptide identification capabilities through the analysis of peptide backbone fragments. Glyco-Decipher has an advantage in identification numbers due to its spectrum expansion method.

Our analysis highlights the trade-offs in glycoproteomics software tools regarding identification sensitivity, consistency, and quantification accuracy. pGlyco3 is the most consistent but less sensitive, MSFragger-Glyco is highly sensitive but prone to false positives, Glyco-decipher achieves the highest identification numbers through spectrum expansion, and StrucGP excels in structural characterization. These findings underscore the need for careful software selection depending on the specific objectives of a glycoproteomics study, particularly in glycan-focused research where quantification consistency remains a challenge.

**Supplementary Note 3. Large-scale and high-quality glycoproteomic spectra enhance prediction performance in existing AI models for N-glycopeptide tandem spectra and retention time.**

The glycoproteomic spectra library and dataset produced in this study not only significantly expanded the scale of the existing database, but also classified the glycopeptide spectra and their identification results into different quality/confidence levels according to the consistency of identification results of different search software. The spectra with consistent identification results obtained by multiple search software are relatively of higher quality, and their identification results are more reliable. This study provides not only a larger amount of glycopeptide data but also high-quality glycopeptide spectra and high-confidence identification results, which can partially make up for the current lack of glycopeptide standard libraries and provide reliable, high-quality training data for AI model training.

We first evaluated whether the different confidence datasets generated in this study could enhance the prediction performance of the DeepGP model^7^. Since the search results reported by pGlyco3, which provide possible glycan structural annotations (PlausibleStruct), were compatible with DeepGP, we selected pGlyco3 subsets and partitioned them based on confidence levels for model training. We trained the model with datasets of ambiguous, low, moderate, 90% of high-confidence data (high_train), as well as a combination of moderate and high_train, with remaining 10% of high-confidence data reserved for testing. We ensured that no glycopeptides overlapped between the training and testing sets. The results demonstrated that data with moderate confidence and above significantly improved the cosine similarity between the predicted and experimental mass spectra, with the median for high_train + moderate reaching 0.95, exceeding previously reported performance (Figure 2e). In contrast, training with low and ambiguous datasets resulted in lower performance, with median cosine similarity of only 0.88 and 0.89, respectively. The performance of the moderate-confidence dataset is comparable to that of the high-confidence dataset, likely due to its larger size (60k vs. 26k glycopeptides) and relatively high data quality, as it was co-identified by two or three different search engines. In addition, the model trained on moderate+high_train data, in the absence of RT calibration, achieved higher Pearson correlation coefficients (PCC) of iRT prediction than the previously reported performance (Figure 2f). This comparison shows that both the size and quality of the training dataset have important impacts on the training effect of the model.

We also evaluated our data using DeepGlyco model, performing 5-fold cross-validation on the high-confidence dataset, where four parts were used for training and one part for validation in each iteration (Figure 2g)^8^. The median dot product (DP) for the whole glycopeptides was consistently above 0.986, and exceeded 0.992 for the glycan parts across all subsets. Furthermore, the spectral angle loss (SA) for glycopeptides ranged from 0.102 to 0.108, while for the glycan parts, it ranged from 0.078 to 0.083. The results were stable, indicating superior data quality and representativeness. Next, we trained the model using the low and moderate datasets, testing on the same high-confidence data aligning with shown in Figure S10a. Although the similarity was slightly lower than high-confidence results mentioned above, it still surpassed the previously reported benchmarks, with SA of 0.150 and DP of 0.972 at the whole spectrum level for the holdout set^8^.

Furthermore, we performed cross-validation partitioning the high-confidence dataset based on different enzymatic patterns. Specifically, we trained the model using data from any two enzymes and validated it against the third one. The similarities dropped marginally, yet the performance remained strong (DP of glycopeptide: 0.983-0.992, SA: 0.081-0.116), reflecting the model's generalization ability (Figure S10b).

In summary, we demonstrated that using a larger dataset generated in this study, particularly the high-quality dataset with consistent identifications from multiple search engines, can aid in training AI models. Moderate and above datasets significantly improve the performance of AI models for MS/MS or retention time prediction. We demonstrated that it is essential to classify the spectra based on confidence using multiple software.

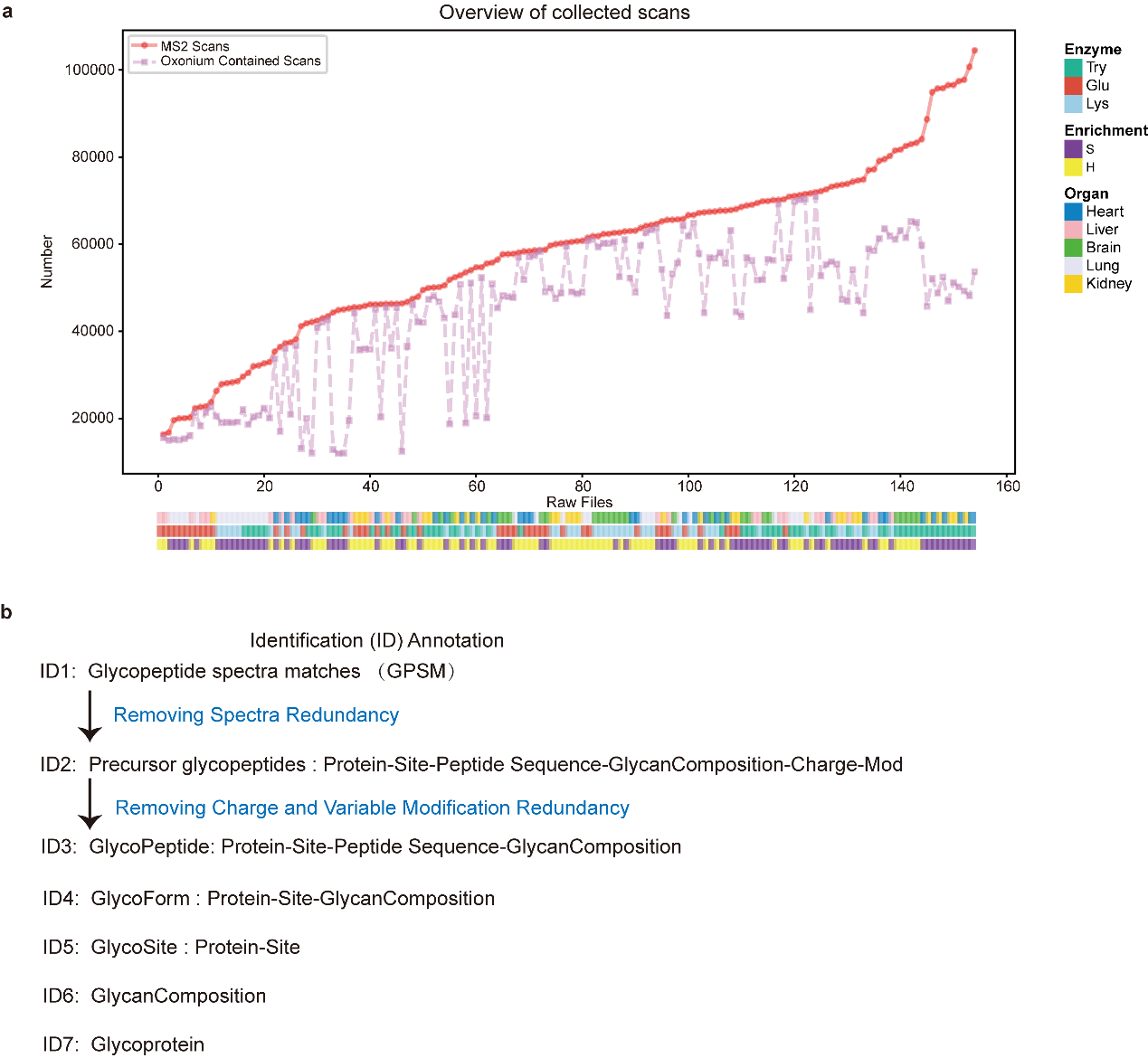

Figure S1. Overview of collected scans and identification annotation process. (a) Overview of collected scans. The plot shows the cumulative number of MS2 scans (red solid line) and oxonium ion-contained scans (purple dashed line) across 154 raw files. Each raw file is represented on the x-axis, while the number of scans is indicated on the y-axis. Color-coded bars beneath the plot represent different experimental conditions, including the enzyme used (Trypsin, Glu-C, Lys-C), enrichment method (Sialic acid enrichment, HILIC), and the organ of origin (Heart, Liver, Brain, Lung, Kidney). (b) Identification (ID) annotation process. A stepwise workflow illustrating the glycopeptide identification and annotation process.

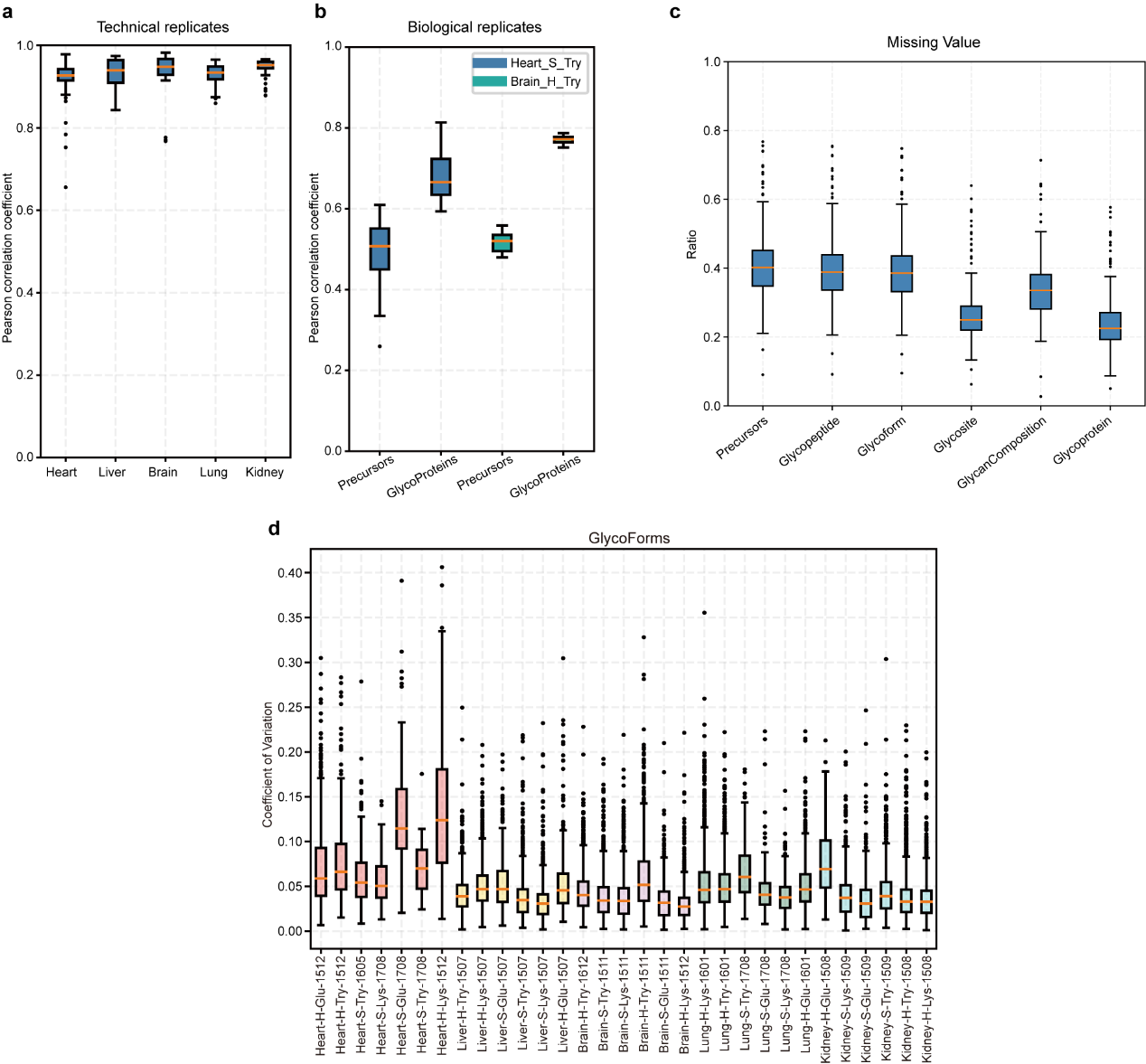

Figure S2. Quality control metrics for N-Glycoproteomic analysis. (a) Pearson correlation coefficients for glycopeptide precursor quantification among technical replicates across different tissues. (b) The reproducibility of glycopeptide precursor and glycoproteins quantification in biological replicates for heart and brain tissues. (c) Analysis of missing values across different IDs (Precursor, Glycopeptide, Glycoform, Glycan, GlycanComposition, Glycoprotein). This analysis helps to assess the completeness and reliability of the glycoproteomic data. (d) The coefficient of variation (CV) for glycoforms across various tissues. Lower CV values indicate higher consistency and precision in glycoprotein quantification. The lower and upper hinges of the box represent the first and third quartiles. The lower and upper whiskers extend from the hinges to the smallest and largest values within 1.5 times the interquartile range (IQR). Source data are provided as a Source Data file.

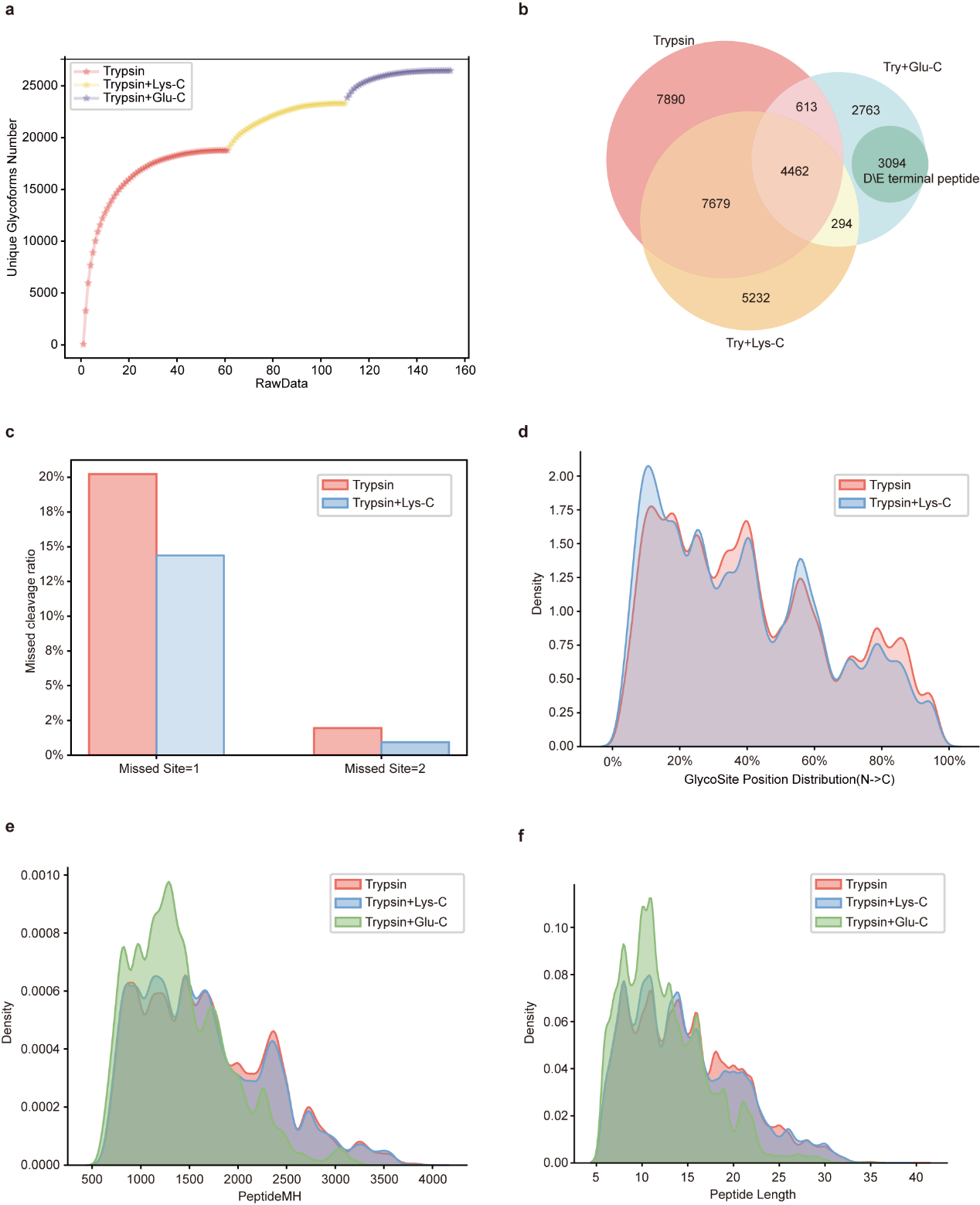

Figure S3: Comparison of glycopeptide analysis using different proteases. (a) Accumulation of unique glycoforms from all raw data for Trypsin, Trypsin+Lys-C, and Trypsin+Glu-C. (b) Venn diagram showing the overlap of glycopeptides identified by Trypsin, Trypsin+Lys-C, and Trypsin+Glu-C. (c) Bar chart showing the ratio of missed cleavage sites for the different proteases. (d) Glycosite position distribution (N→C) for the glycopeptides identified by Trypsin and Trypsin+Lys-C. (e) Peptide mass distribution (PeptideMH) for the glycopeptides identified by each protease. (f) Distribution of peptide lengths for the glycopeptides identified by each protease.

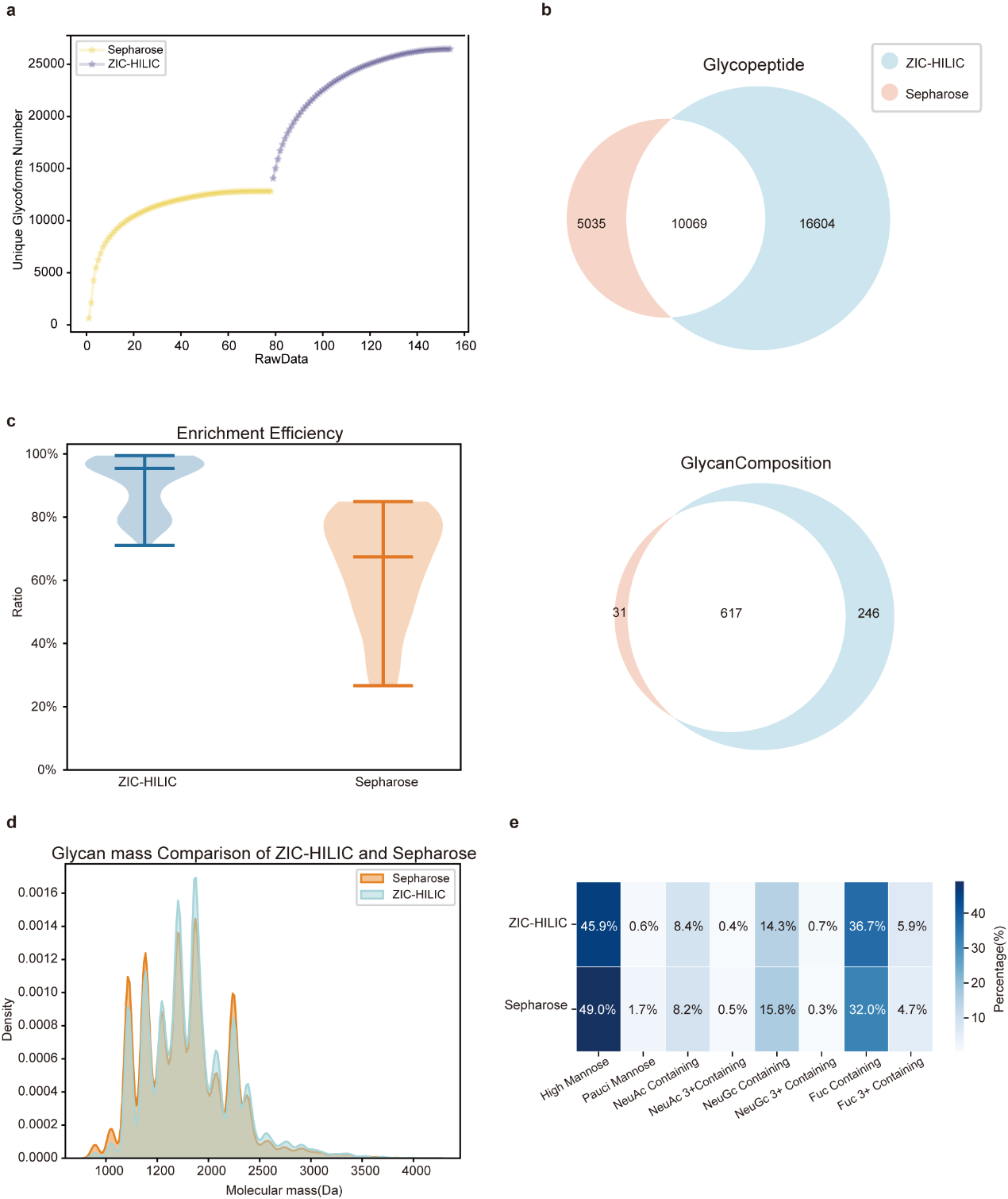

Figure S4. Comparison of glycopeptide and glycan compositions between Sepharose and ZIC-HILIC enrichment methods. (a) Accumulation of unique glycoforms as a function of raw data. ZIC-HILIC and Sepharose exhibit different numbers of unique glycopeptides. (b) Venn diagrams showing the overlap of glycopeptides and glycan compositions identified by ZIC-HILIC and Sepharose. (c) Enrichment efficiency comparison between ZIC-HILIC and Sepharose, with ZIC-HILIC showing a higher overall enrichment ratio. The lower and upper hinges of the box represent the first and third quartiles. The lower and upper whiskers extend from the hinges to the smallest and largest values within 1.5 times the interquartile range (IQR). Source data are provided as a Source Data file. (d) Glycan mass comparison of ZIC-HILIC and Sepharose, displaying the molecular mass distribution of glycans. (e) Heatmap showing the percentage distribution of different glycan types (e.g., High Mannose, NeuAc-containing, Fuc-containing).

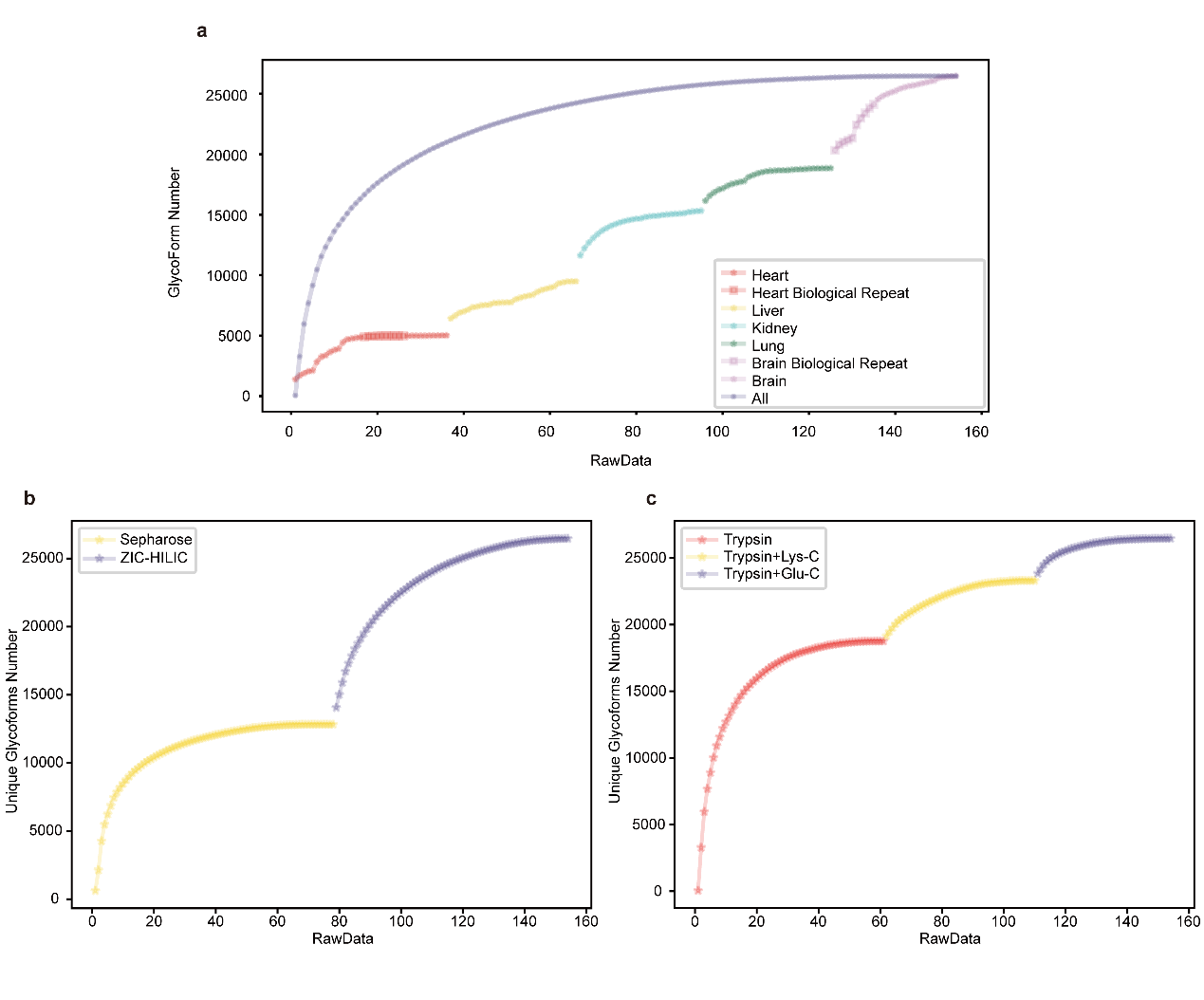

Figure S5. Saturation curves of unique glycoform identification across various tissues.

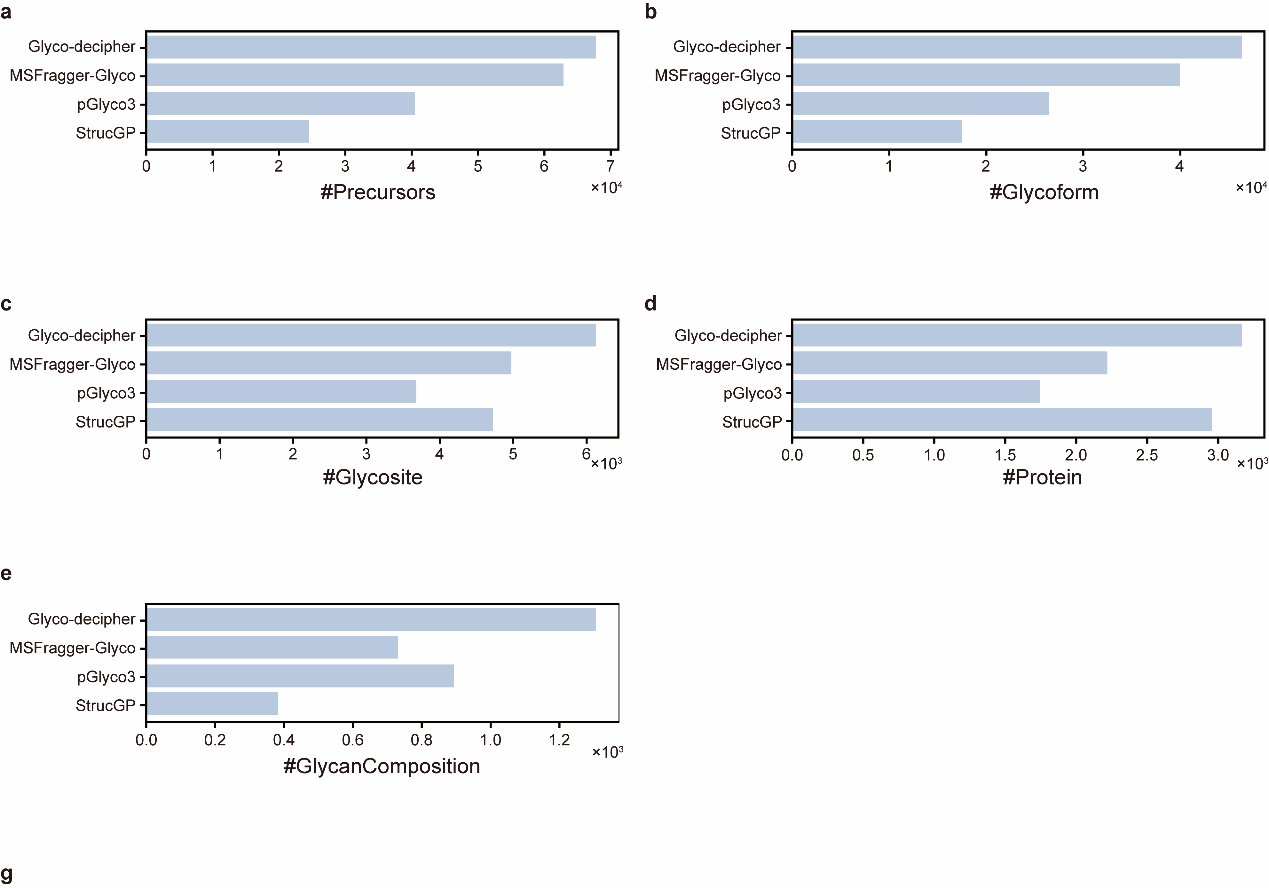

Figure S6: Comprehensive analysis of glycoproteomic data across different software tools. (a-e) The number of identified precursors (a), glycoforms (b), glycosites (c), proteins (d), and glycan compositions (e) using different software tools.

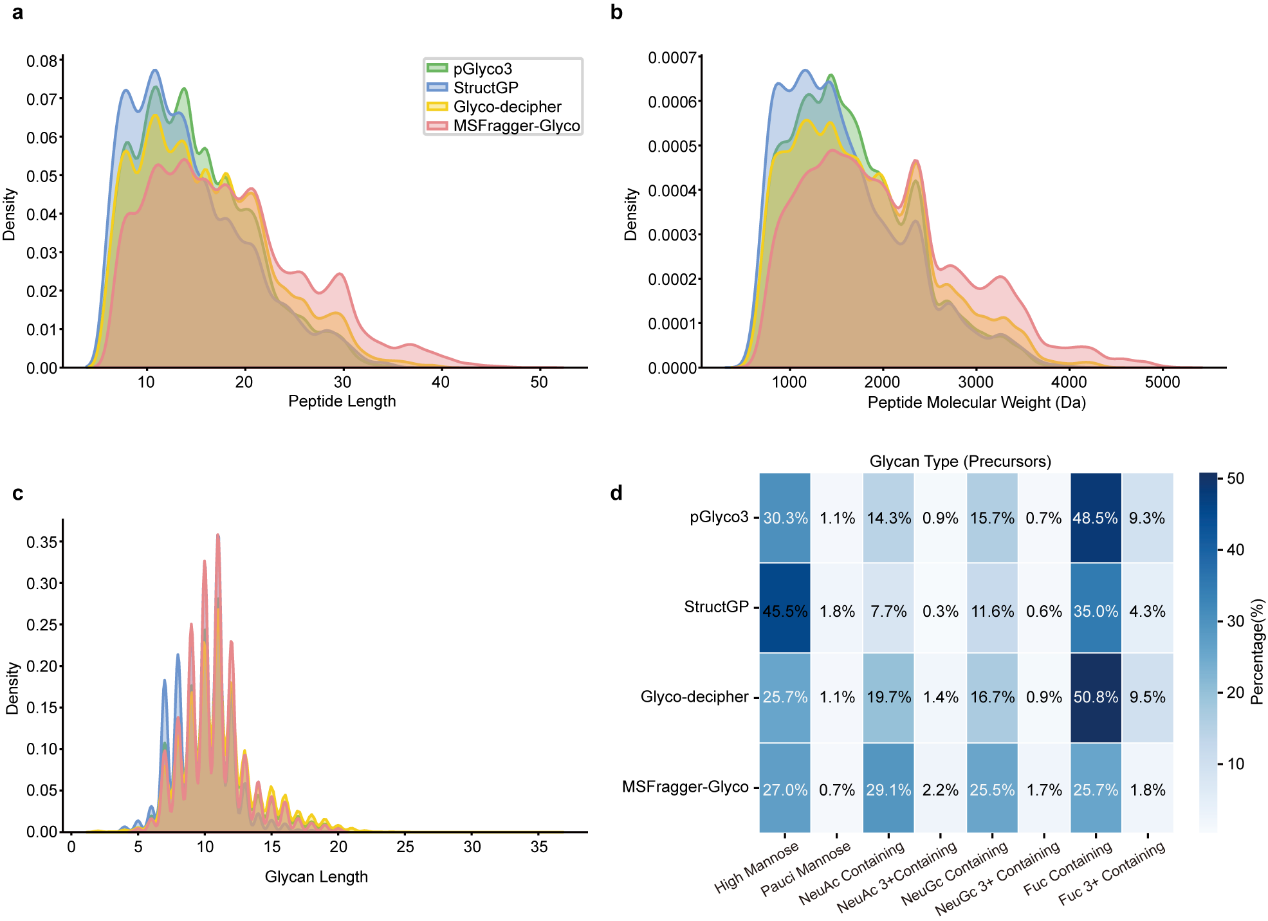

Figure S7. Comparison of glycoproteomics analysis results across different software tools. (a) Density distribution of peptide lengths identified by pGlyco3, StructGP, Glyco-decipher, and MSFragger-Glyco. The x-axis represents peptide length, while the y-axis indicates density. (b) Density distribution of peptide molecular weights (Da) identified by the four tools. (c) Density distribution of glycan lengths identified by the four tools. (d) Proportion of different glycan types (precursors) identified by the four tools. The glycan types are categorized into high mannose, pauci mannose-containing, NeuAc-2+ containing, NeuAc-3+ containing, NeuGc-2+ containing, NeuGc-3+ containing, Fuc-2+ containing, and Fuc-3+ containing. The intensity of the blue color represents the percentage of each glycan type identified by each tool.

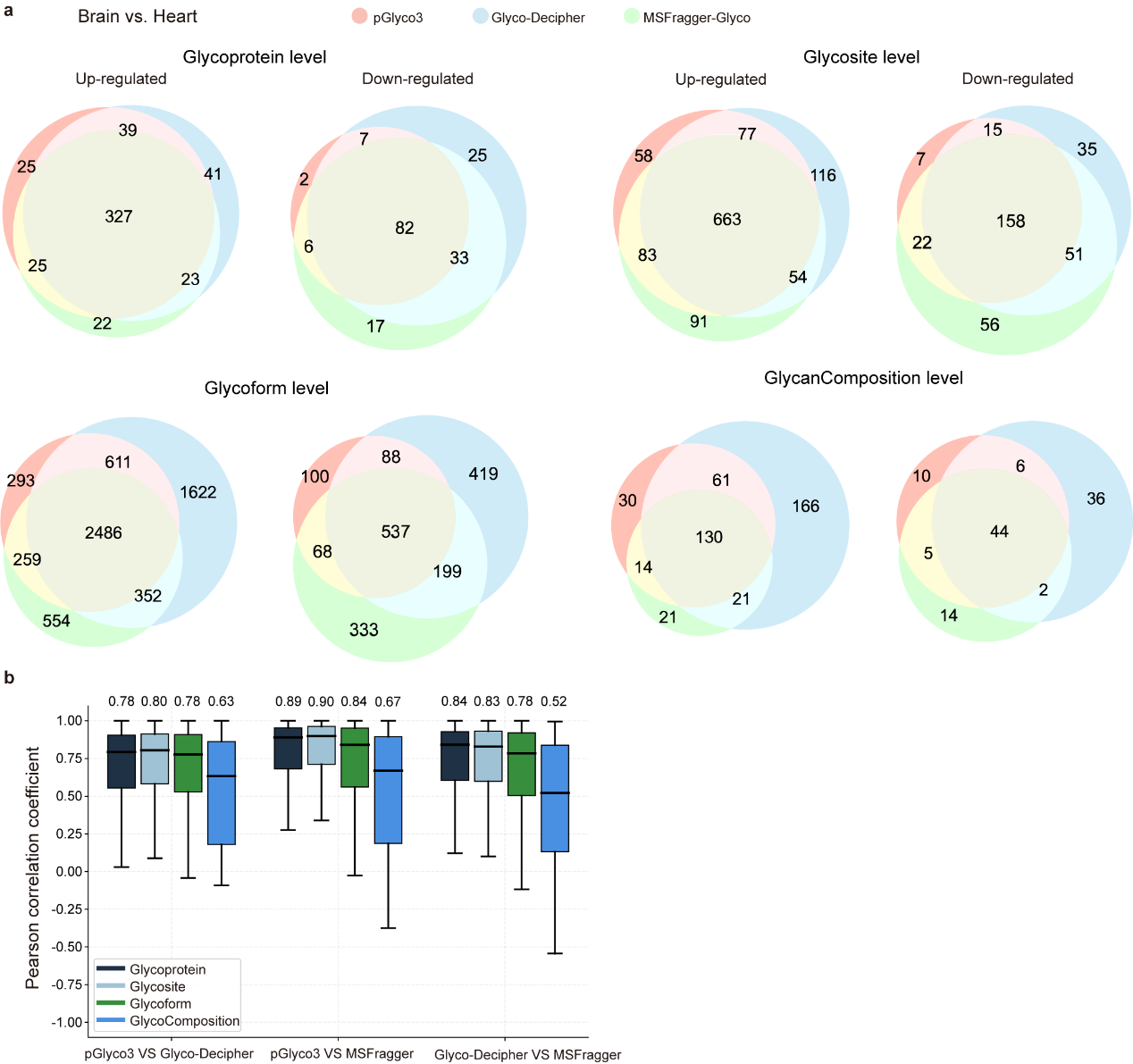

Figure S8. Comparison of Quantitative Glycoproteomics Data Between Brain and Heart Samples Using Different Software with Trypsin and Trypsin+Lys-C Digestion. (a) Venn diagrams showing the overlap of up-regulated and down-regulated glycoproteins, glycoforms, glycosites, and glycan compositions between brain and heart samples, as quantified by three software tools: pGlyco3, Glyco-Decipher, and MSFragger-Glyco. (b) Boxplots of Pearson correlation coefficients comparing the glycoprotein, glycosite, glycoform, and glycan composition data commonly identified and quantified by the three software. The comparisons include pGlyco3 vs. Glyco-Decipher, pGlyco3 vs. MSFragger, and Glyco-Decipher vs. MSFragger. Higher correlation values indicate better agreement in the quantification values between the platforms. Source data are provided as a Source Data file. The lower and upper hinges of the box represent the first and third quartiles. The lower and upper whiskers extend from the hinges to the smallest and largest values within 1.5 times the interquartile range (IQR). Source data are provided as a Source Data file.

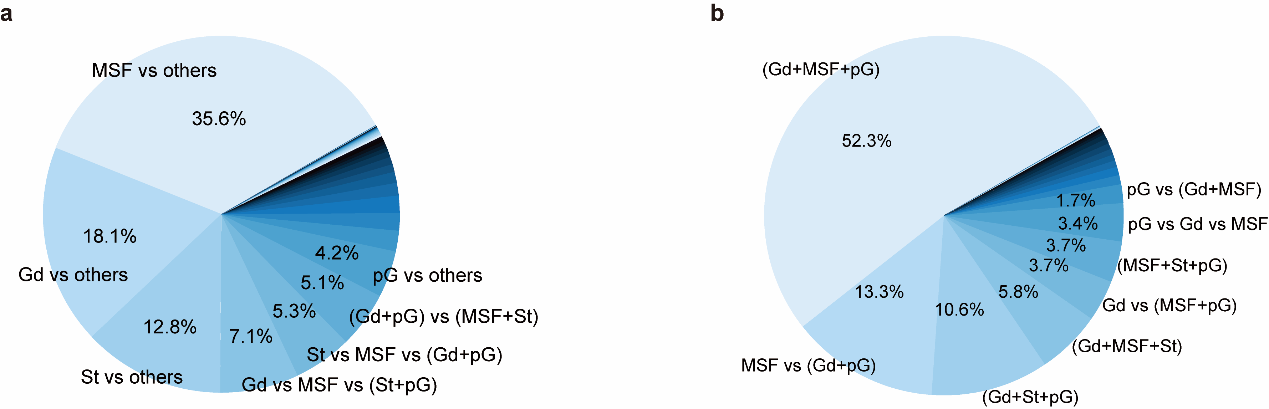

Figure S9. Analysis of inconsistent precursor identifications across four software tools. (a) The pie chart illustrates the distribution of spectra with inconsistent precursor identifications among four glycoproteomics software tools: MSFragger-Glyco (MSF), pGlyco3 (pG), Glyco-decipher (Gd), and StrucGP (St). It highlights spectra where one tool reports a unique identification or where groups of tools produce distinct results, excluding the cases where all four tools agree. For example, "MSF vs others" indicates spectra where MSFragger-Glyco reported a unique result compared to the other tools, which agreed on a different precursor. (b) The pie chart shows the precursor identification analysis for spectra identified by any three of the four tools. Each section represents the percentage of spectra with matching or differing precursor identifications. For example, "Gd+MSF+pG" represents spectra where Glyco-decipher, MSFragger-Glyco, and pGlyco3 all identified the same precursor, while "MSF vs (Gd + pG)" shows spectra where MSFragger-Glyco identified a different precursor compared to Glyco-decipher and pGlyco3, which agreed on the same one.

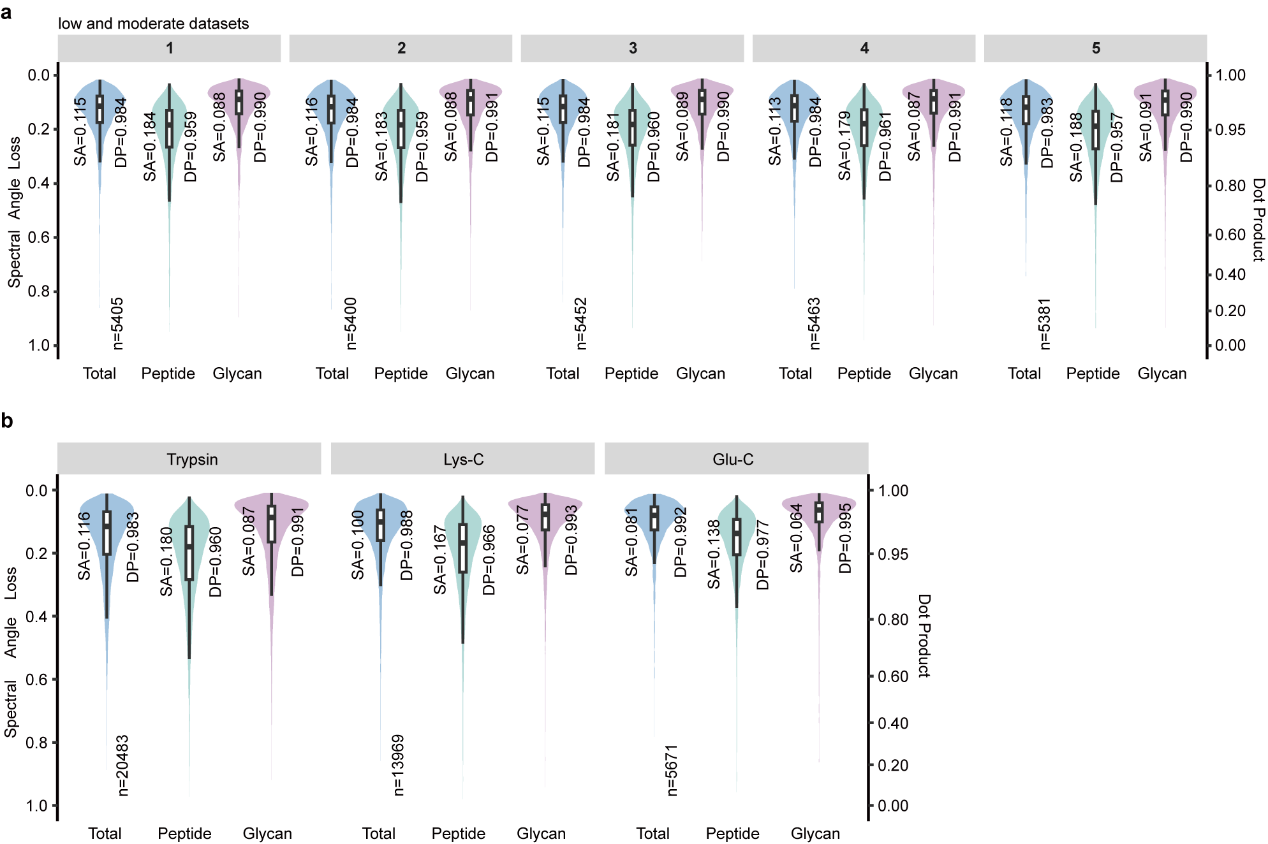

Figure S10. Enhancement of AI training performance with datasets of varying quality. (a) Distribution of spectral similarities using DeepGlyco trained on the low and moderate datasets and tested on the same high-conf subset as in Fig 2g. (e) Distribution of spectral similarities in cross-validation, with the high-conf dataset partitioned according to different enzymatic patterns. The lower and upper hinges of the box represent the first and third quartiles. The lower and upper whiskers extend from the hinges to the smallest and largest values within 1.5 times the interquartile range (IQR). Source data are provided as a Source Data file.

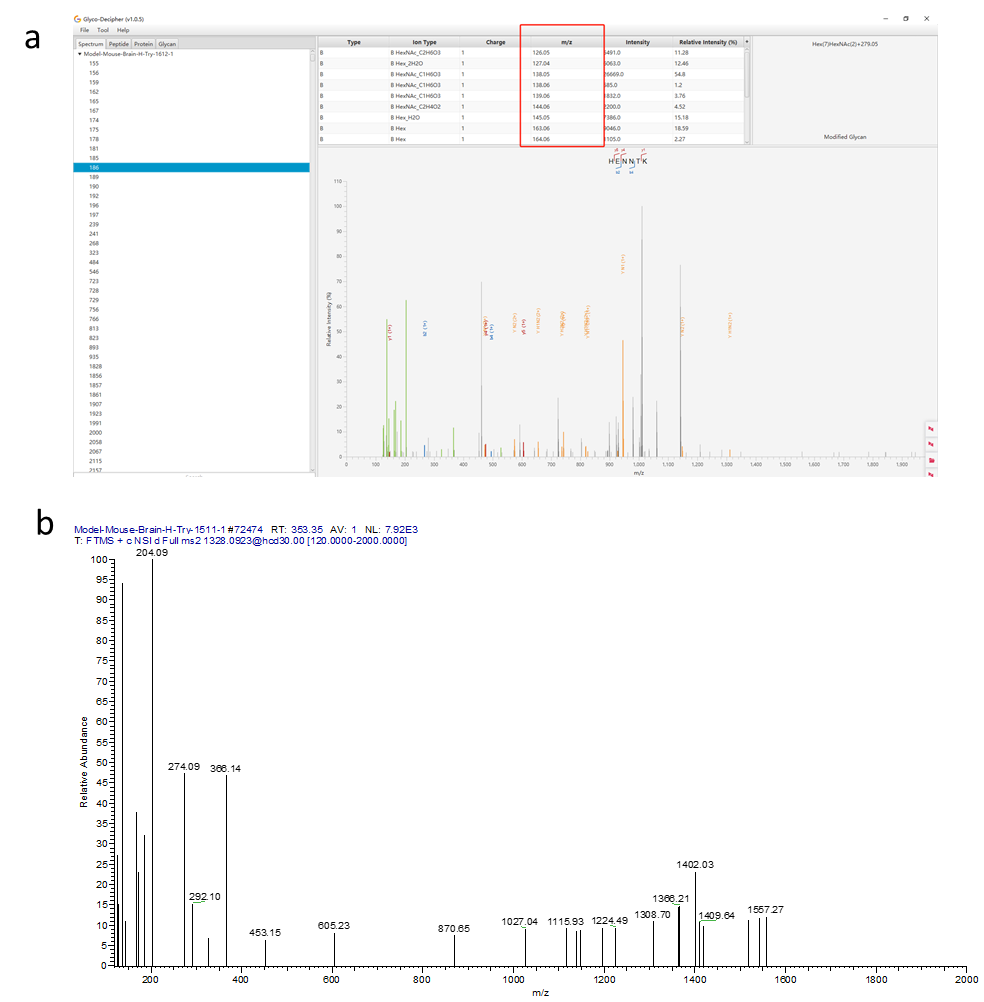

Figure S11. Example spectra for unidentified spectra. (a) A low-quality spectrum showing limited ion information, which makes it difficult to accurately identify the glycopeptide. (b) An example spectrum with abundant ion information, but the glycopeptide was not identified by any software.

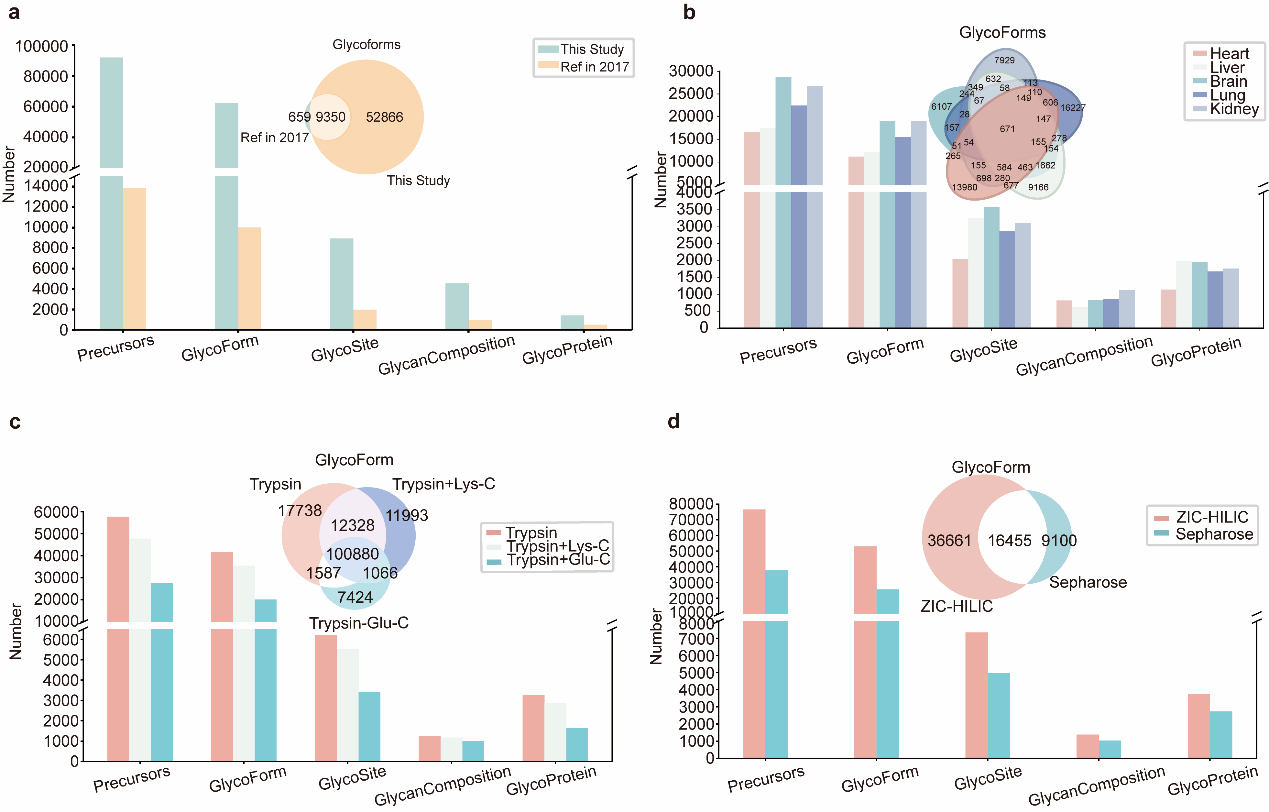

Figure S12. Comparison and Identification of Glycoforms and Glycoproteins Across Mouse Tissues and Enzyme Treatments. (a) Comparison of precursor, glycoform, glycosite, glycan composition, and glycoprotein identifications between this study and a reference study from 2017. (b) Bar chart depicting the number of glycoforms identified in different mouse tissues and a Venn diagram showing the overlap of glycoforms across these tissues. (c) The number of identified precursors, glycoforms, glycosites, glycan compositions, and glycoproteins for different enzyme treatments. The Venn diagram within this panel shows the overlap of glycoform identifications. (d) The number of identified glycoforms, glycosites, glycan compositions, and glycoproteins between two enrichment methods (ZIC-HILIC and Sepharose).

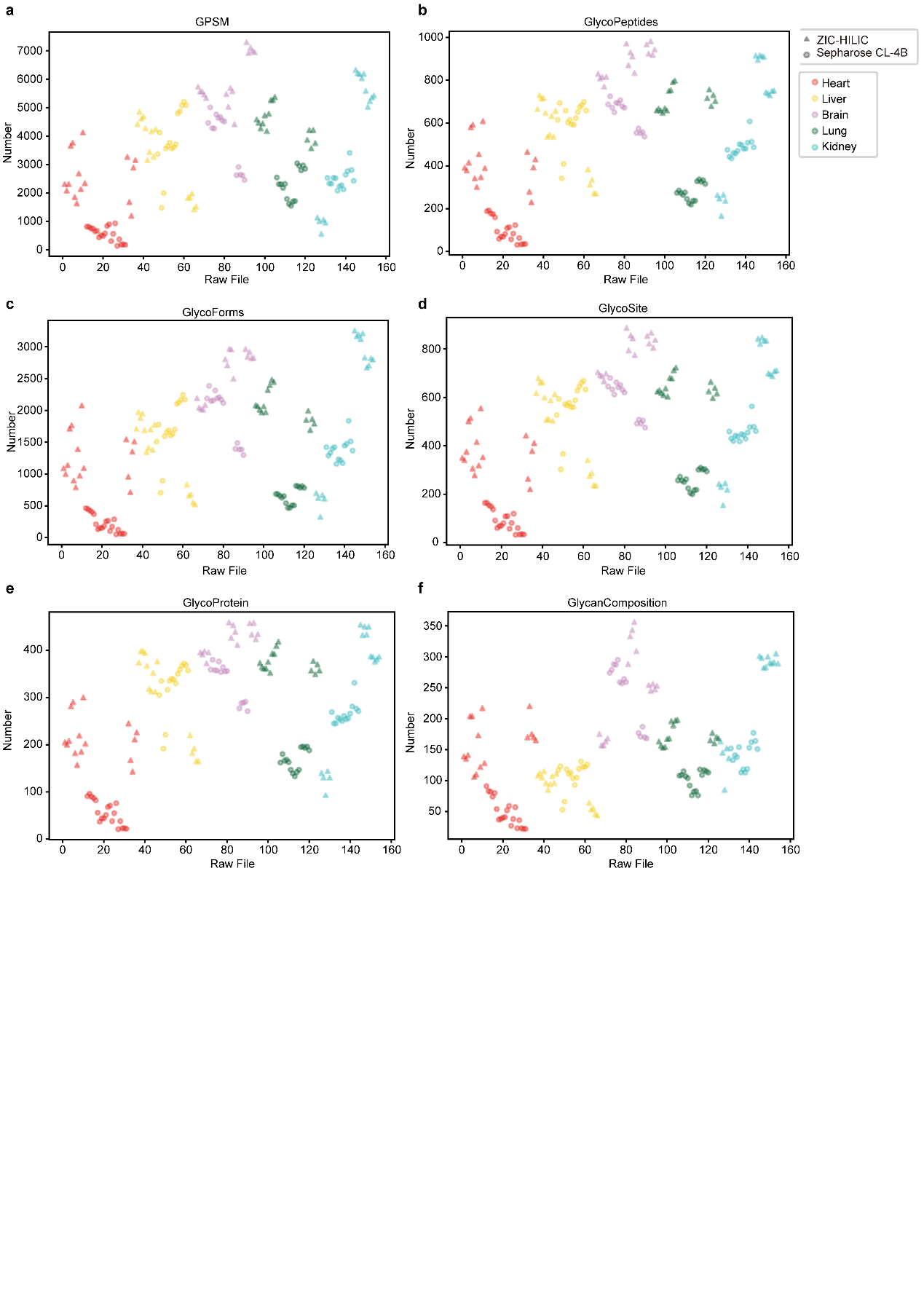

Figure S13: Comparison of Glycoproteomic Metrics(pGlyco3 results) Across Different Tissues and Enrichment Methods. The number of GPSMs (a), glycopeptides (b), glycoforms (c), glycosite (d), glycoproteins (e) and glycan composition (f) identified in each raw file across the different tissues. Each point represents a raw file, with colors and shapes indicating the tissue and enrichment method, respectively.

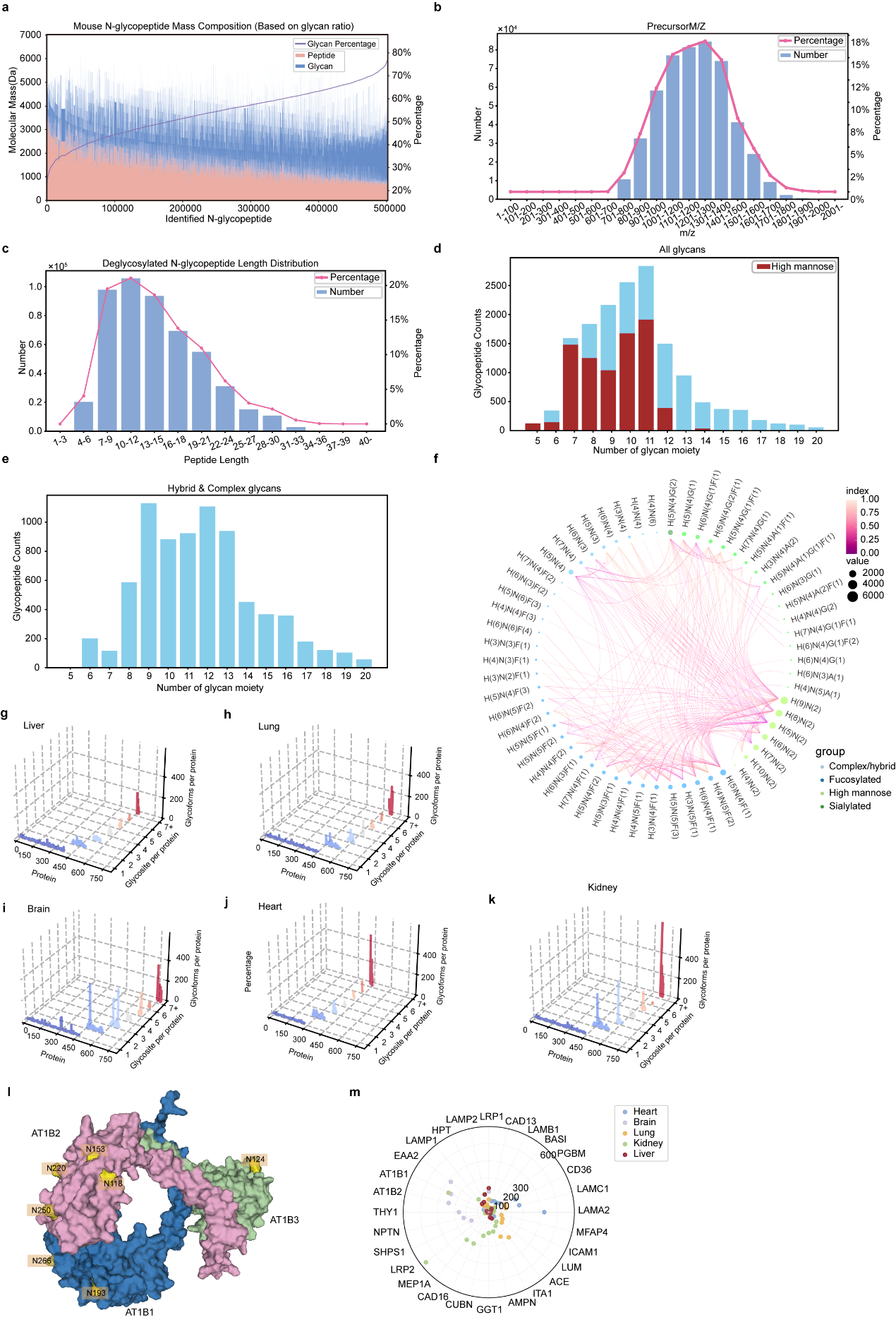

Figure S14: Analysis of Glycoproteomic Data Across Multiple Tissues and Structural Insights. (a) illustrates the mass composition of mouse N-glycopeptides based on glycan ratio. The histogram shows the distribution of glycan mass percentages relative to peptide mass across identified N-glycopeptides, highlighting the diversity of glycan structures. (b) shows the distribution of precursor m/z (mass-to-charge ratio) values for identified glycopeptides. The histogram illustrates the range and frequency of m/z values, indicating the ionization efficiency and mass distribution of the glycopeptides. (c) presents the length distribution of deglycosylated N-glycopeptides. The bar chart shows the number and percentage of peptides of different lengths, providing insights into the peptide characteristics post-deglycosylation. (d) Glycan composition analysis showing the glycopeptide counts for glycans with varying numbers of glycan moieties. High mannose glycan types are highlighted in red. (e) Distribution of glycopeptide counts with different numbers of hybrid and complex type glycan moieties. (f) Network representation of different glycan structures categorized by type (complex/hybrid, fucosylated, high mannose, sialylated) with node size representing the number of occurrences and color indicating different glycan types. An edge is drawn between two nodes if the corresponding glycan types co-occur at the same site. (g-k) present 3D scatter plots showing the distribution of glycopeptides identified in the Liver (g), lung (h), Brain (i), Heart (j), and Kidney (k). The axes represent the protein index (X-axis), the number of glycosylation sites per protein (Y-axis), and the total number of glycoforms associated with each protein (Z-axis). Each plot highlights tissue-specific glycoproteomic profiles. (i) shows the structural model of a representative glycoprotein complex (AT1B1, AT1B2, AT1B3) with identified glycosylation sites highlighted. The structural insights provide a visual representation of how glycosylation might influence protein function and interactions. (m) displays a radar plot comparing the abundance of glycoproteins across different tissues. Key glycoproteins are marked.

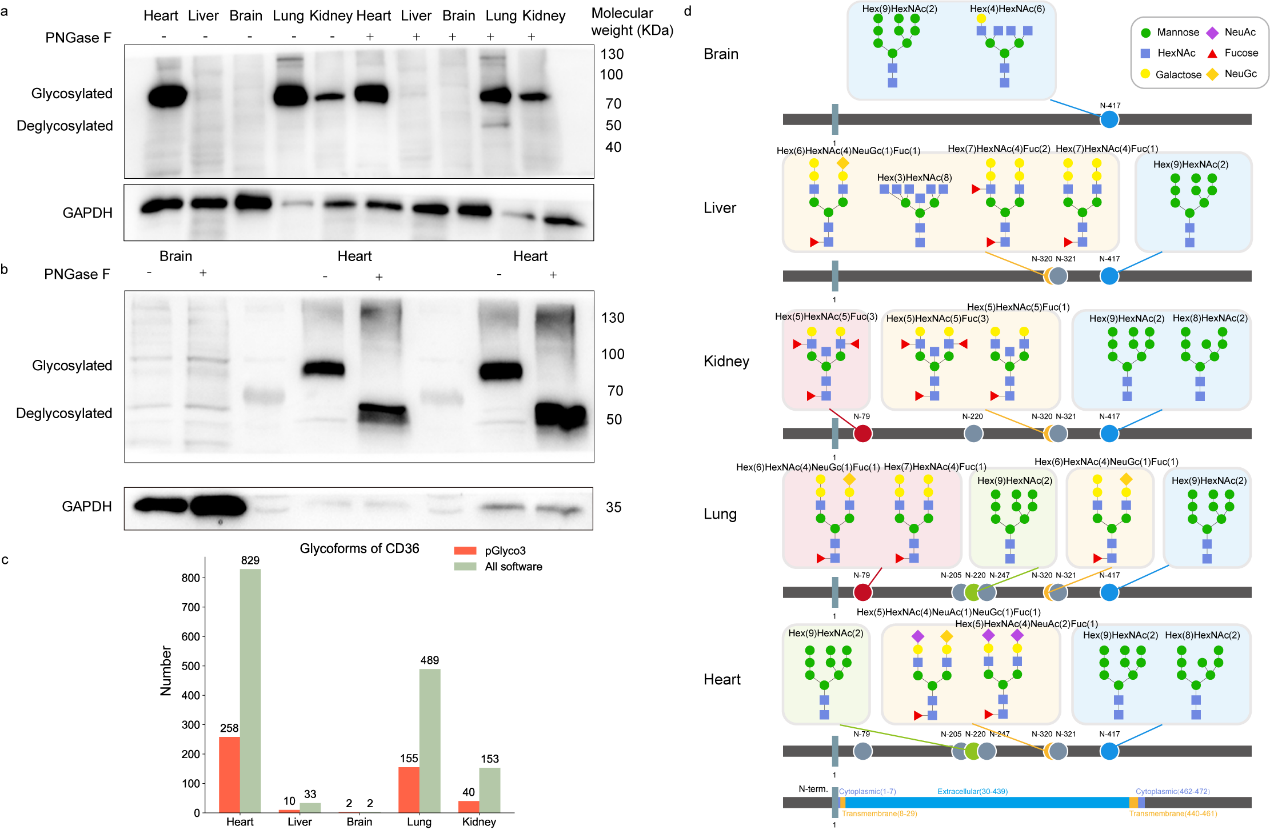

Figure S15. Western blot analysis and glycoform identification of CD36 glycoprotein across different tissues. (a) Western blot analysis of CD36 glycoprotein under native conditions, demonstrating the presence of intact glycoprotein in various tissues. Under native conditions, PNGase F treatment did not remove the N-glycans from the CD36, indicating the intact glycosylation state of the protein. (b) Western blot analysis of CD36 glycoprotein under denatured conditions, where PNGase F treatment was applied to assess the effect of glycan removal on protein migration. The removal of N-linked glycans by PNGase F resulted in a shift in the molecular weight of the CD36 protein, highlighting the impact of glycosylation on the protein's electrophoretic behavior. (c) Comparison of glycoforms numbers of CD36 identified in different tissues. (d) The top five CD36 glycoforms, ranked by their intensities in pGlyco3, are shown across various tissues. The glycan composition and possible structures are displayed, illustrating the diversity of glycoforms in different tissue types.

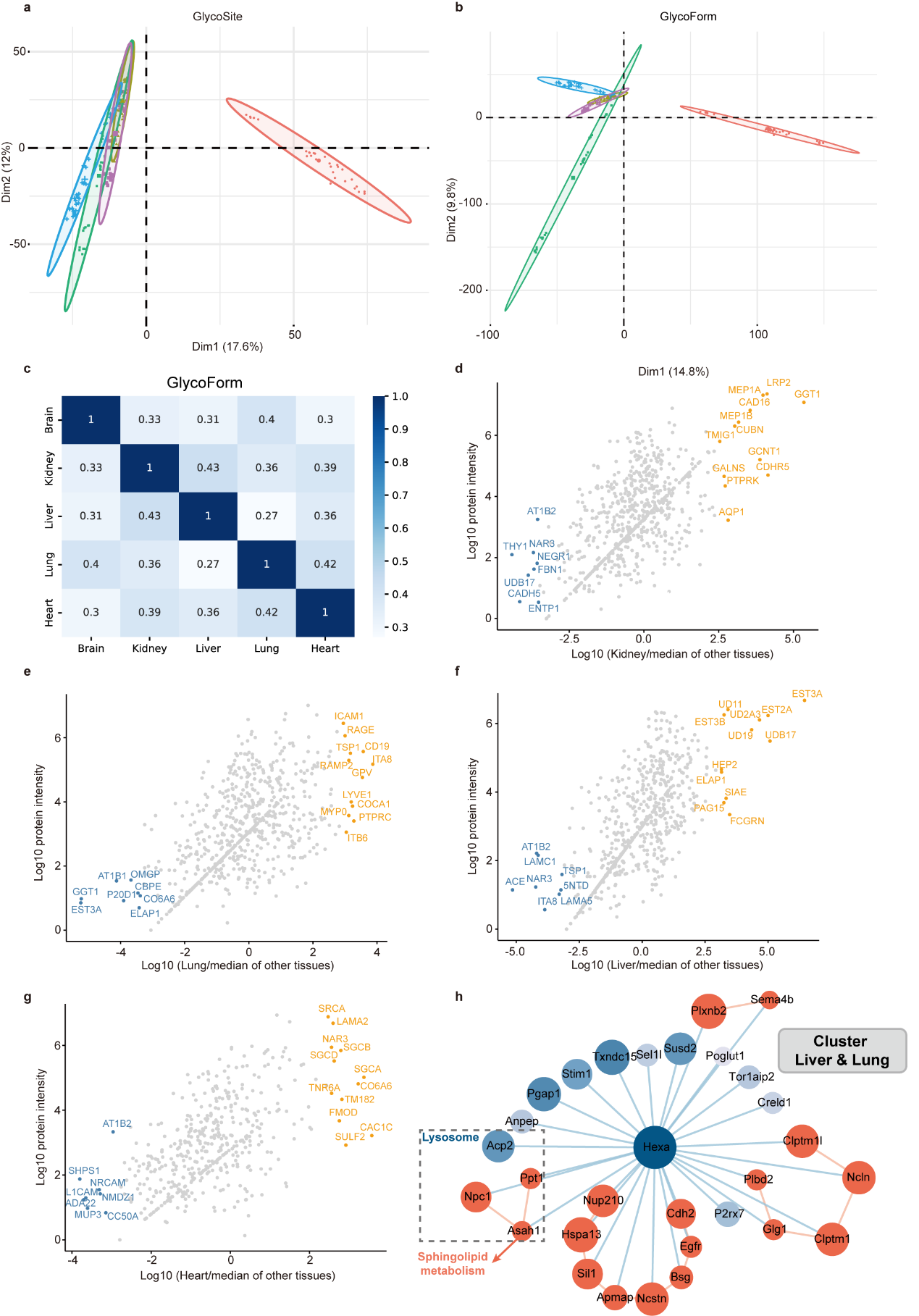

Figure S16: Comparative Analysis of Glycosylation Patterns Across Different Tissues. (a) shows a PCA plot of glycosylation sites (GlycoSite) identified across different tissues. Each color represents a different tissue, and the clustering indicates tissue-specific glycosylation patterns. (b) displays a PCA plot of glycoforms, further highlighting distinct glycosylation profiles among the tissues. (c) presents a heatmap showing the correlation of glycoform intensities between different tissues. Higher correlation values indicate similar glycosylation patterns, while lower values suggest tissue-specific glycosylation. (d-g) provide scatter plots comparing the log10 intensities of glycoproteins between each tissue and the others. (h) illustrates a network analysis focusing on glycoproteins with enriched functions in Liver and Lung. The network highlights clusters of proteins involved in lysosome and sphingolipid metabolism, with nodes representing glycoproteins and edges indicating functional relationships.

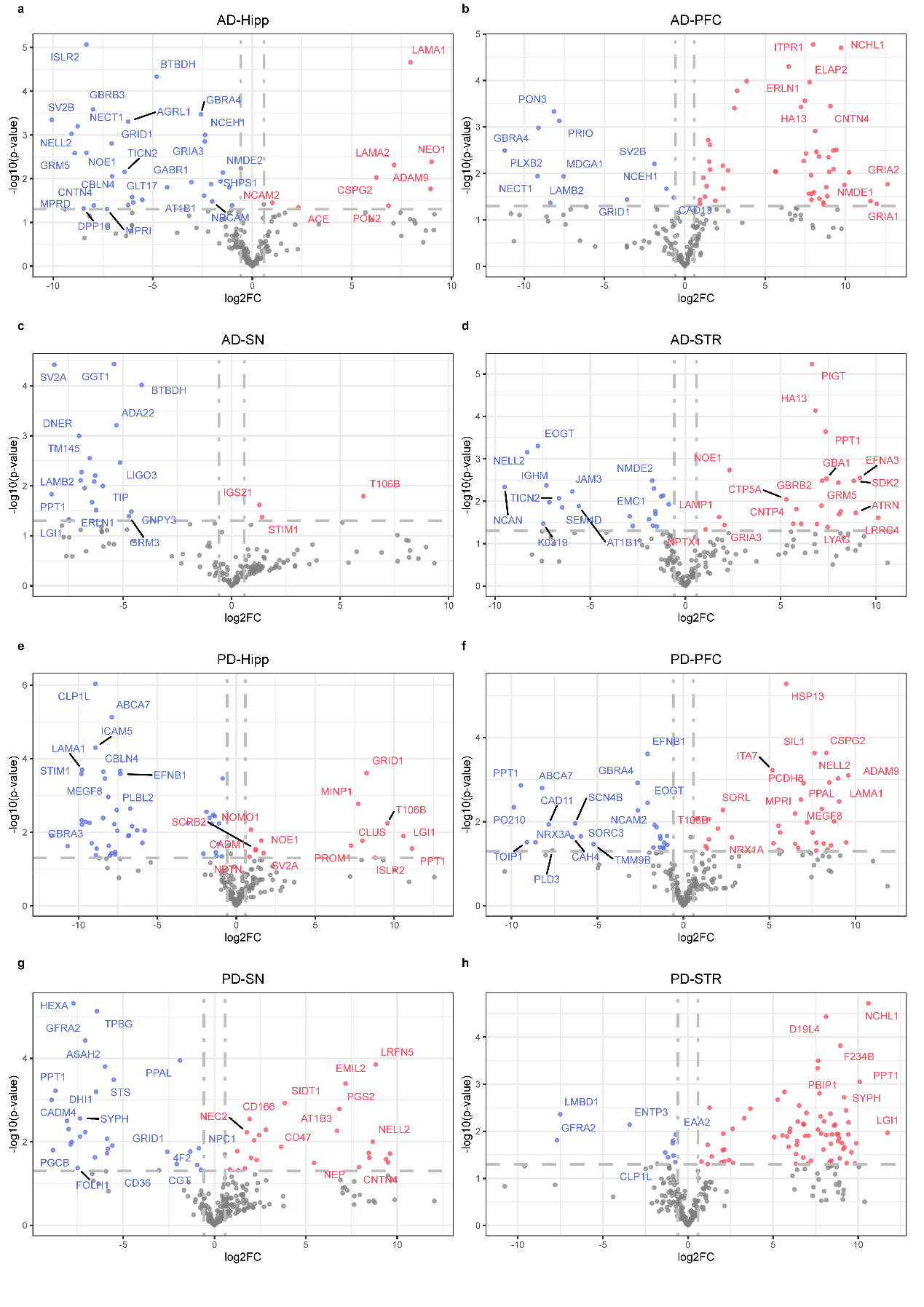

Figure S17. Volcano plots showing differential protein expression in various brain regions of Alzheimer's disease (AD) and Parkinson's disease (PD) patients. Each plot represents a specific brain region: (a) Hippocampus (AD-Hipp), (b) Prefrontal Cortex (AD-PFC), (c) Substantia Nigra (AD-SN), (d) Striatum (AD-STR), (e) Hippocampus (PD-Hipp), (f) Prefrontal Cortex (PD-PFC), (g) Substantia Nigra (PD-SN), and (h) Striatum (PD-STR). Red dots represent upregulated proteins, and blue dots represent downregulated proteins, based on log2 fold change (log2FC) and -log10 p-value. Gray dots indicate non-significant proteins. The vertical dashed lines indicate the threshold for significant fold changes. Statistical significance was determined using a two-tailed independent two-sample t-test to compare group means. P-values < 0.05 were considered significant. Source data are provided as a Source Data file.

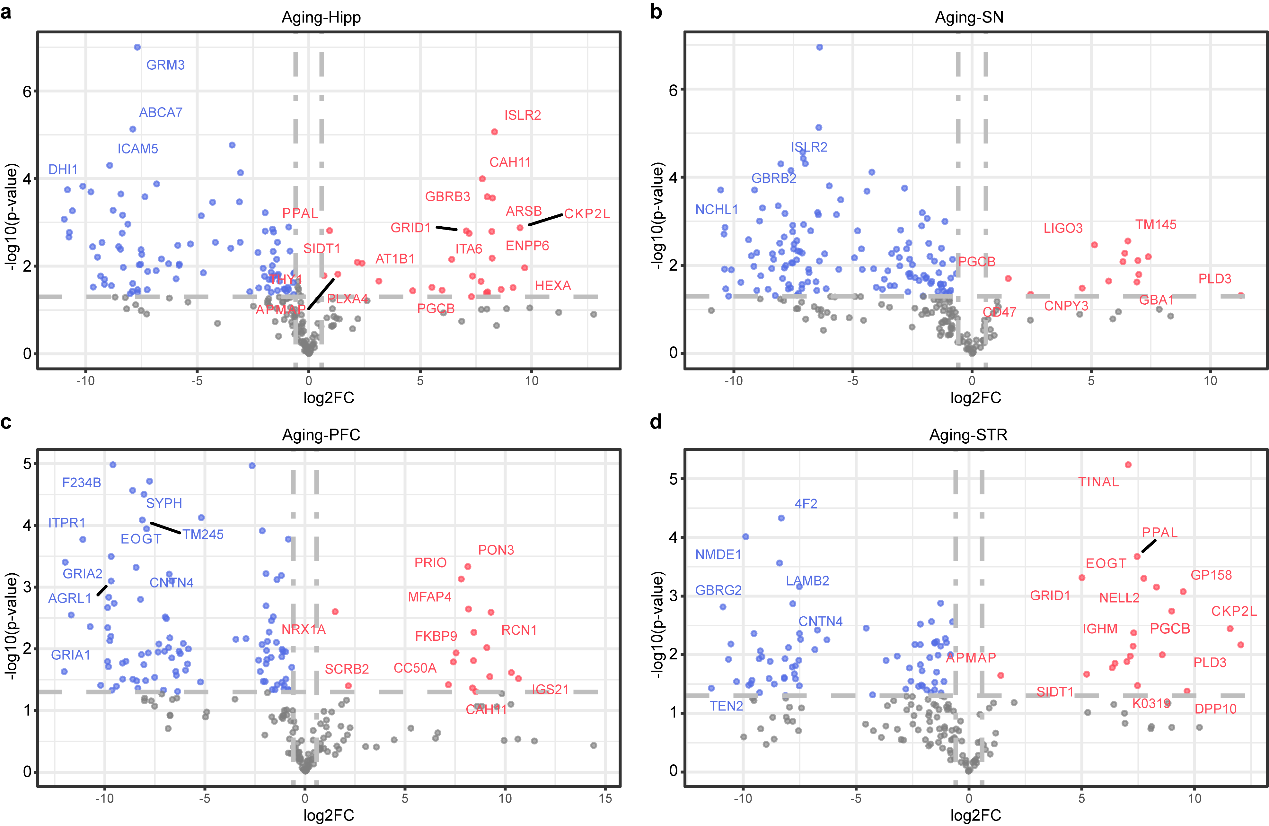

Figure S18. Volcano plots depicting differential protein expression associated with aging in different brain regions. Each plot represents a specific brain region: (a) Hippocampus (Aging-Hipp), (b) Substantia Nigra (Aging-SN), (c) Prefrontal Cortex (Aging-PFC), and (d) Striatum (Aging-STR). Red dots indicate upregulated proteins, and blue dots indicate downregulated proteins based on log2 fold change (log2FC) and -log10 p-value. Gray dots represent non-significant proteins. The vertical dashed lines represent the threshold for significant fold changes. The plots highlight proteins significantly altered during the aging process in each region. Statistical significance was determined using a two-tailed independent two-sample t-test to compare group means. P-values < 0.05 were considered significant. Source data are provided as a Source Data file.

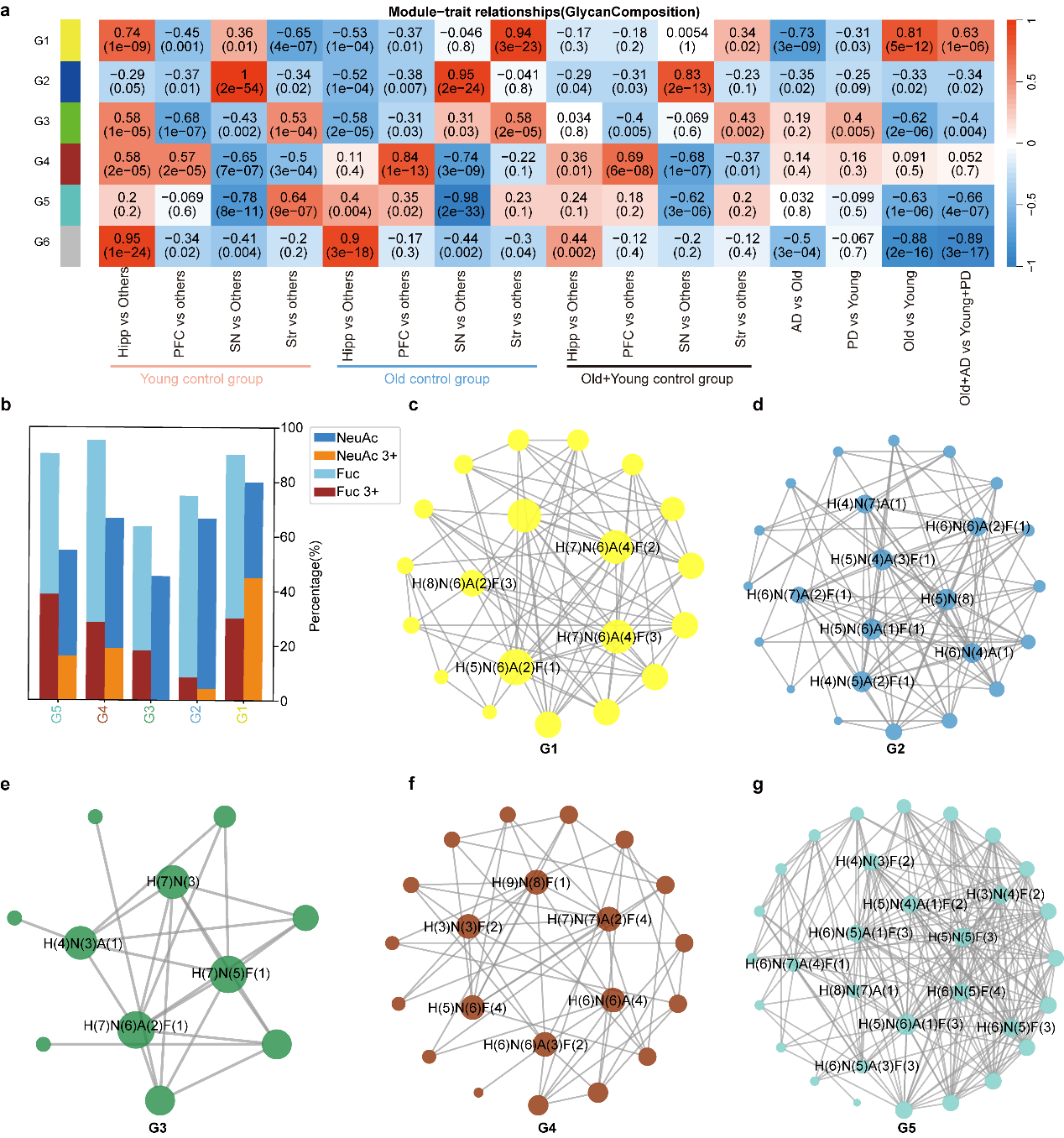

Figure S19. Module-Trait Relationships for Glycan Composition Analysis. (a) Heatmap showing the module-trait relationships for different glycan compositions across various brain regions (Hippocampus, PFC, SN, STR) and control groups (Young, Old, and Combined). The color scale represents correlation coefficients, with red indicating positive correlation and blue indicating negative correlation; p-values are shown in parentheses. (b) Bar graph displaying the percentage composition of different glycans (NeuAc, NeuAc 3+, Fuc, Fuc 3+) in modules G1 to G6.
(c-g) Network plots for glycan modules G1 to G6, illustrating the connectivity and relationships between key glycans, with nodes representing individual glycans and edges representing their interactions. Different colors indicate distinct glycan modules. Source data are provided as a Source Data file.

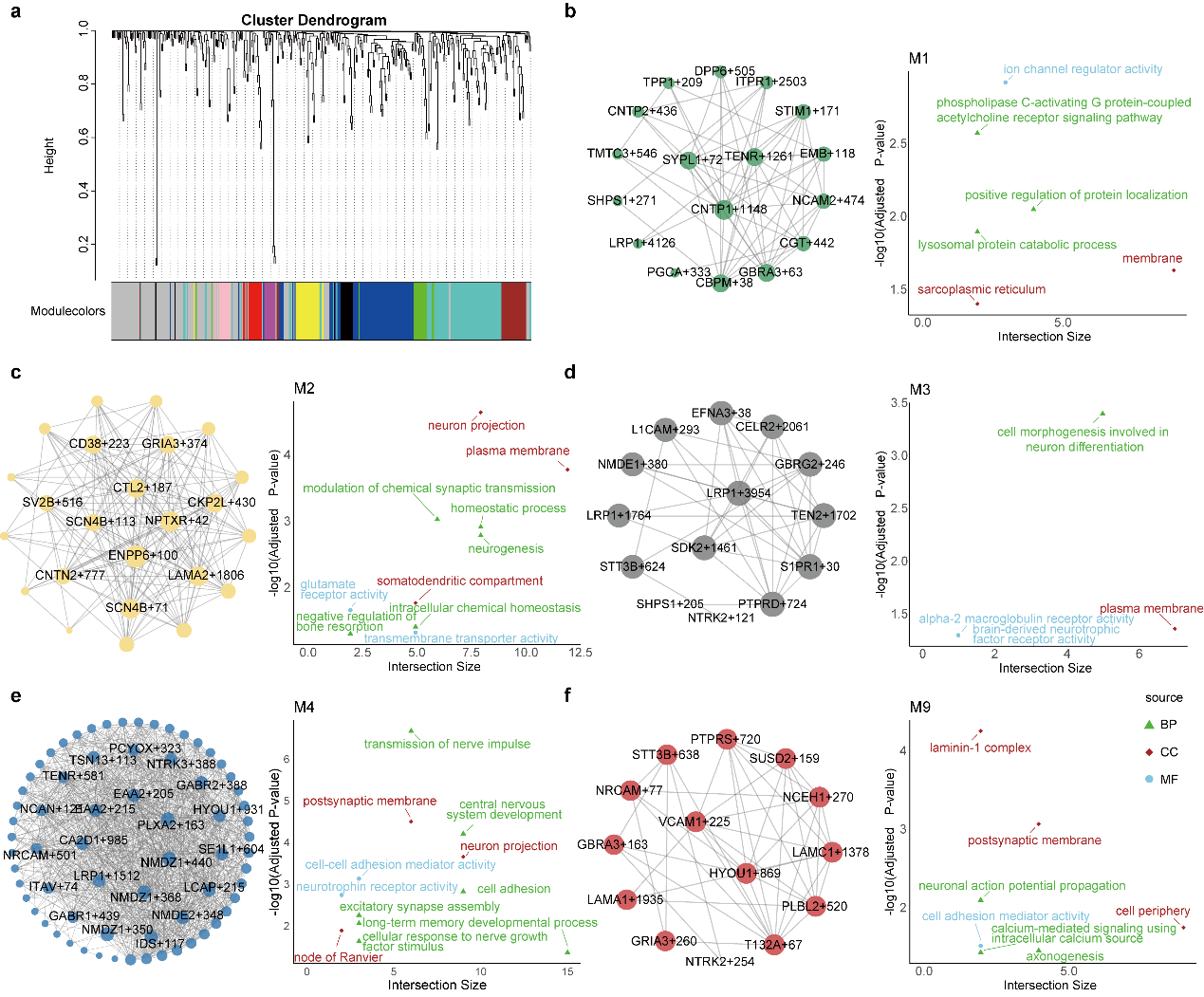

Figure S20. Hierarchical Clustering and Functional Enrichment Analysis of Glycosite Modules. (a) Cluster dendrogram showing hierarchical clustering of glycosites into modules based on their expression profiles, with different colors representing distinct glycosite modules. (b-f) Network diagrams and associated enriched functional terms for different modules identified in the clustering:(b) Module M1: Enriched in processes such as ion channel regulator activity and positive regulation of protein localization. (c) Module M2: Related to modulation of chemical synaptic transmission and neurogenesis. (d) Module M3: Involved in cell morphogenesis in neuron differentiation. (e) Module M4: Associated with transmission of nerve impulses and postsynaptic membrane functions. (f) Module M9: Includes terms related to laminin-1 complex and neuronal action potential propagation. Gene Ontology (GO) terms are shown with colors representing biological processes (BP), cellular components (CC), and molecular functions (MF). P-values were calculated using a hypergeometric test and adjusted for multiple testing using the Benjamini–Hochberg FDR method. Terms with adjusted p-values < 0.05 were considered significant. Source data are provided as a Source Data file.

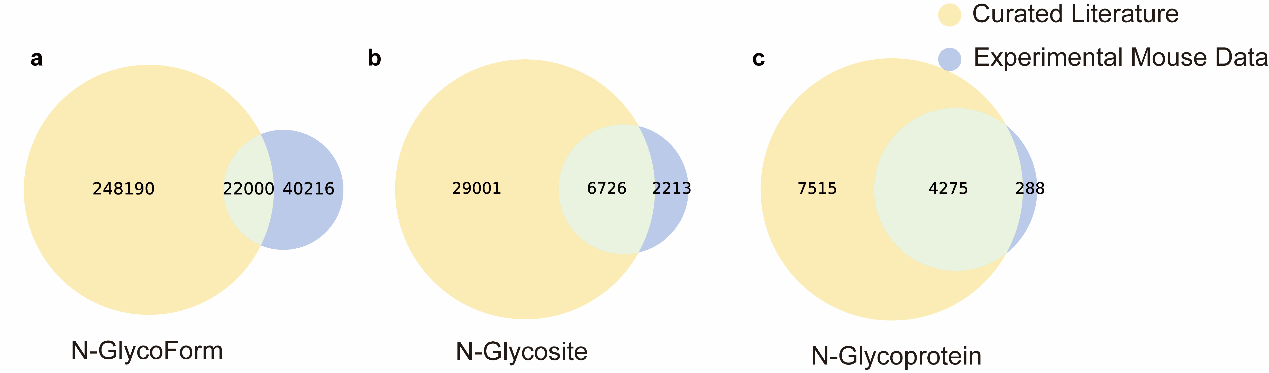

Figure S21. Venn diagrams illustrating the overlap between curated literature and experimental mouse data for three categories: (a) N-GlycoForms, (b) N-Glycosites, and (c) N-Glycoproteins.

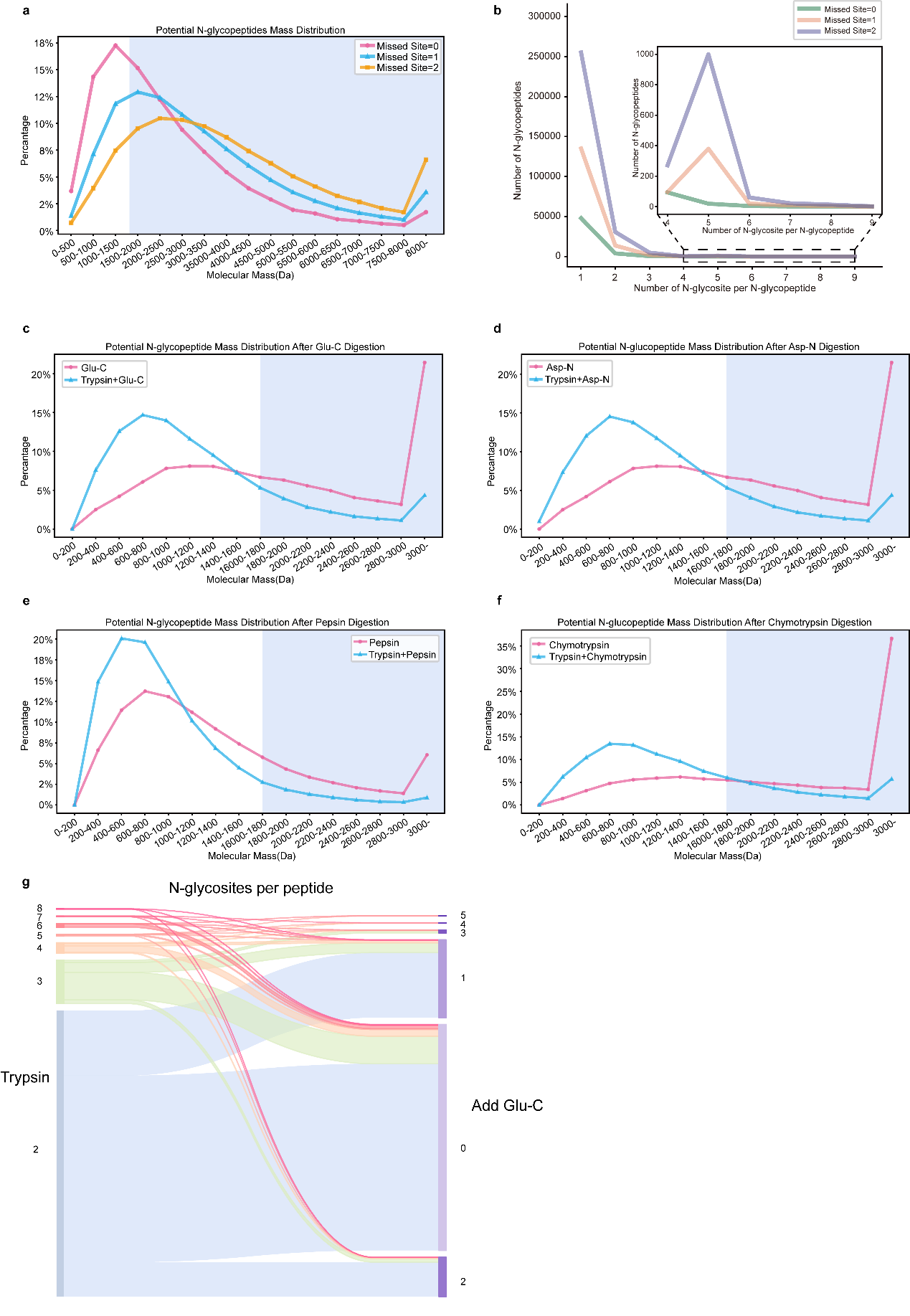

Figure S22. Analysis of N-Glycopeptide Mass Distribution and Glycosite Reduction Across Different Enzyme Digestions. (a) Distribution of potential N-glycopeptide masses after trypsin digestion, grouped by the number of missed cleavage sites (0, 1, or 2). The molecular masses are displayed on the x-axis (in Da), and the percentage of N-glycopeptides is shown on the y-axis. (b) Number of N-glycosites per N-glycopeptide after trypsin digestion, stratified by the number of missed cleavage sites (0, 1, or 2). The main plot shows the number of N-glycosites (x-axis) versus the number of N-glycopeptides (y-axis), with an inset providing a closer look at peptides containing 4 to 9 glycosites. (c-f) Mass distribution of potential N-glycopeptides after digestion with different enzymes. Each panel compares the distribution between single-enzyme and combined enzyme digestions:(c) Glu-C alone versus Trypsin + Glu-C; (d) Asp-N alone versus Trypsin + Asp-N; (e) Pepsin alone versus Trypsin + Pepsin; (f) Chymotrypsin alone versus Trypsin + Chymotrypsin. The x-axis represents the molecular mass (in Da) of N-glycopeptides, while the y-axis shows their percentage. (g) Sankey plot depicting the reduction in the number of N-glycosites per peptide after Glu-C digestion following initial trypsin digestion. The flow lines indicate the changes in the number of glycosites (y-axis) before (left) and after (right) the addition of Glu-C.

**Source Data:**

Figure S15a

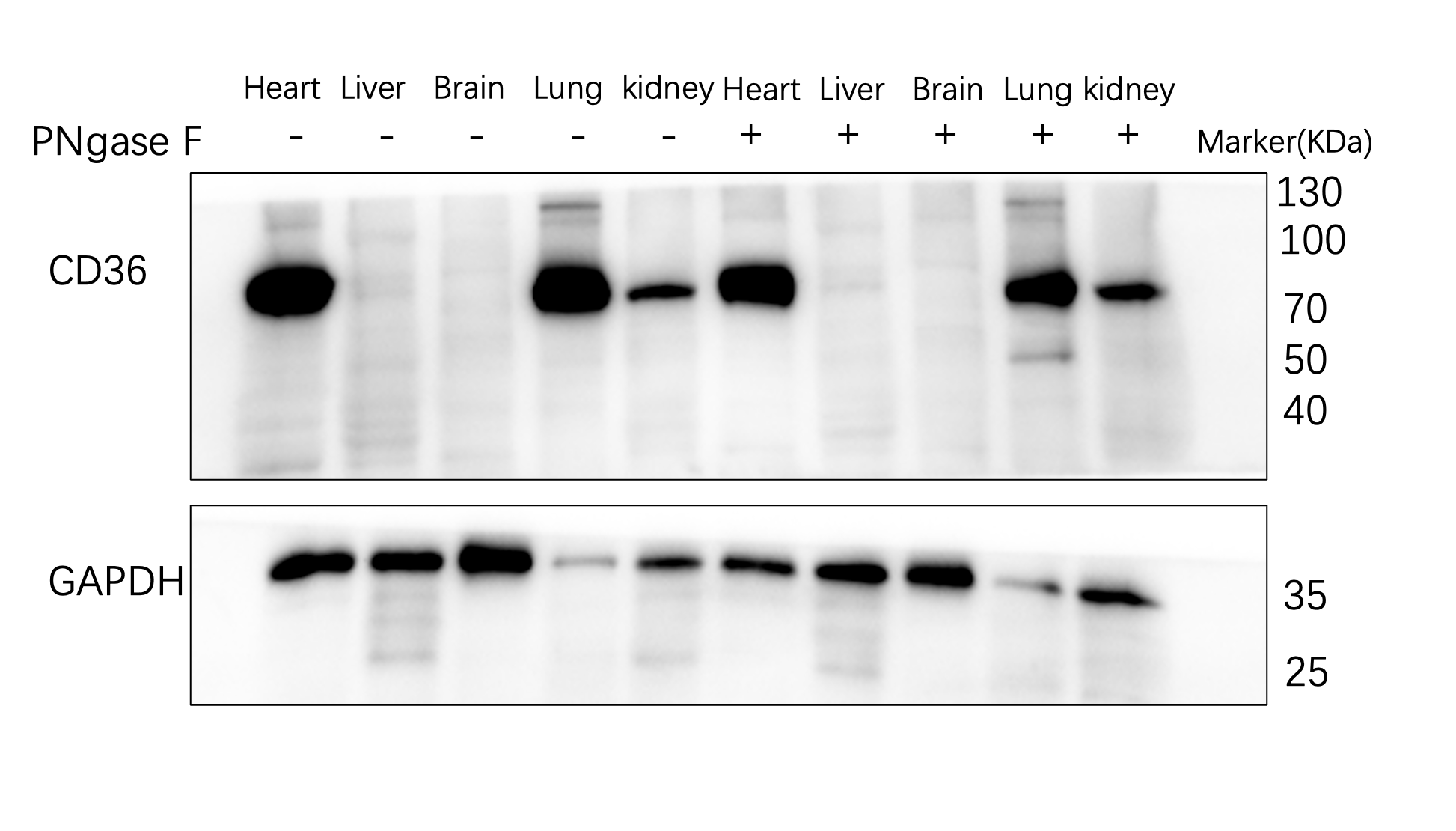

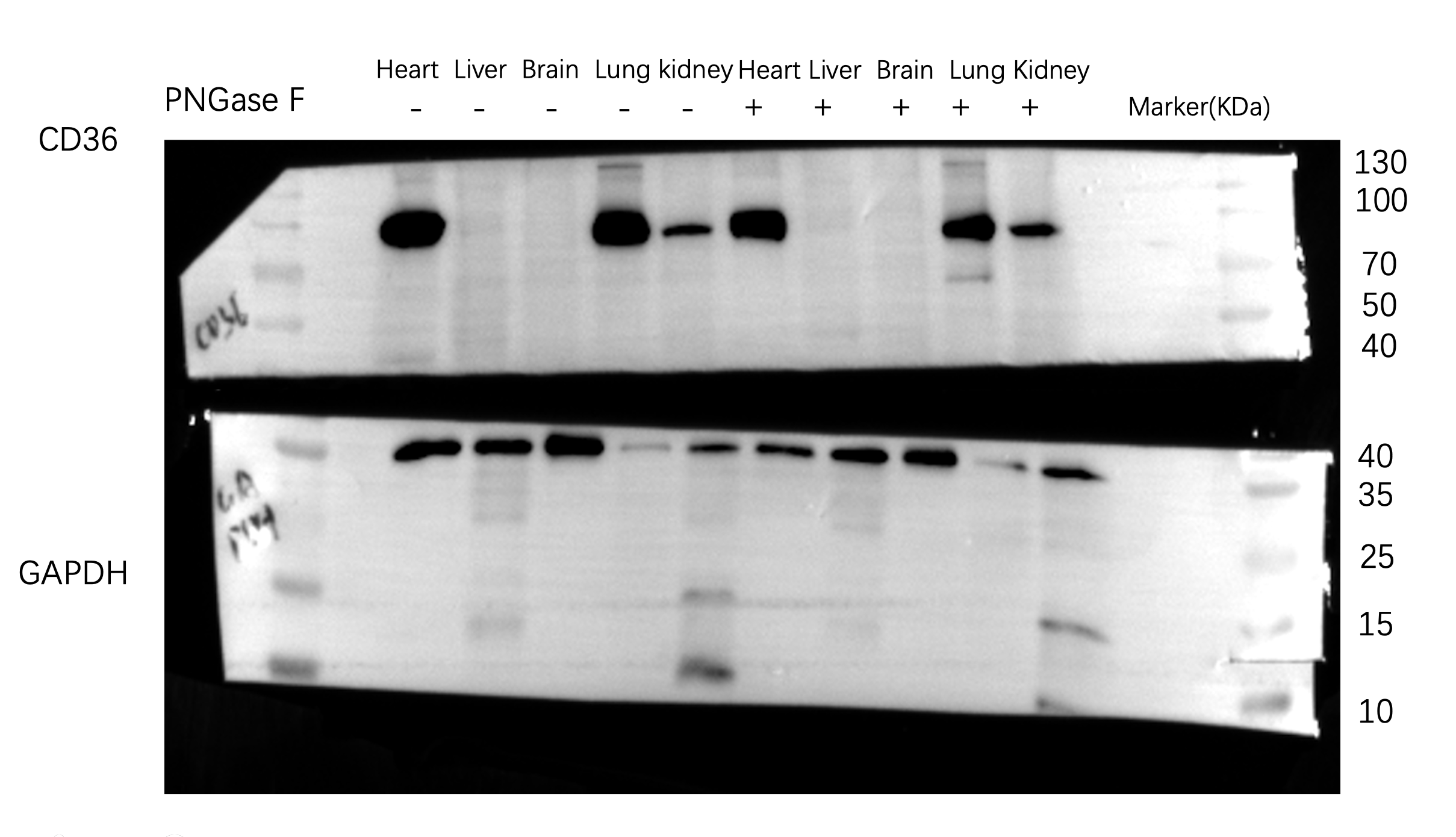

Figure S15b

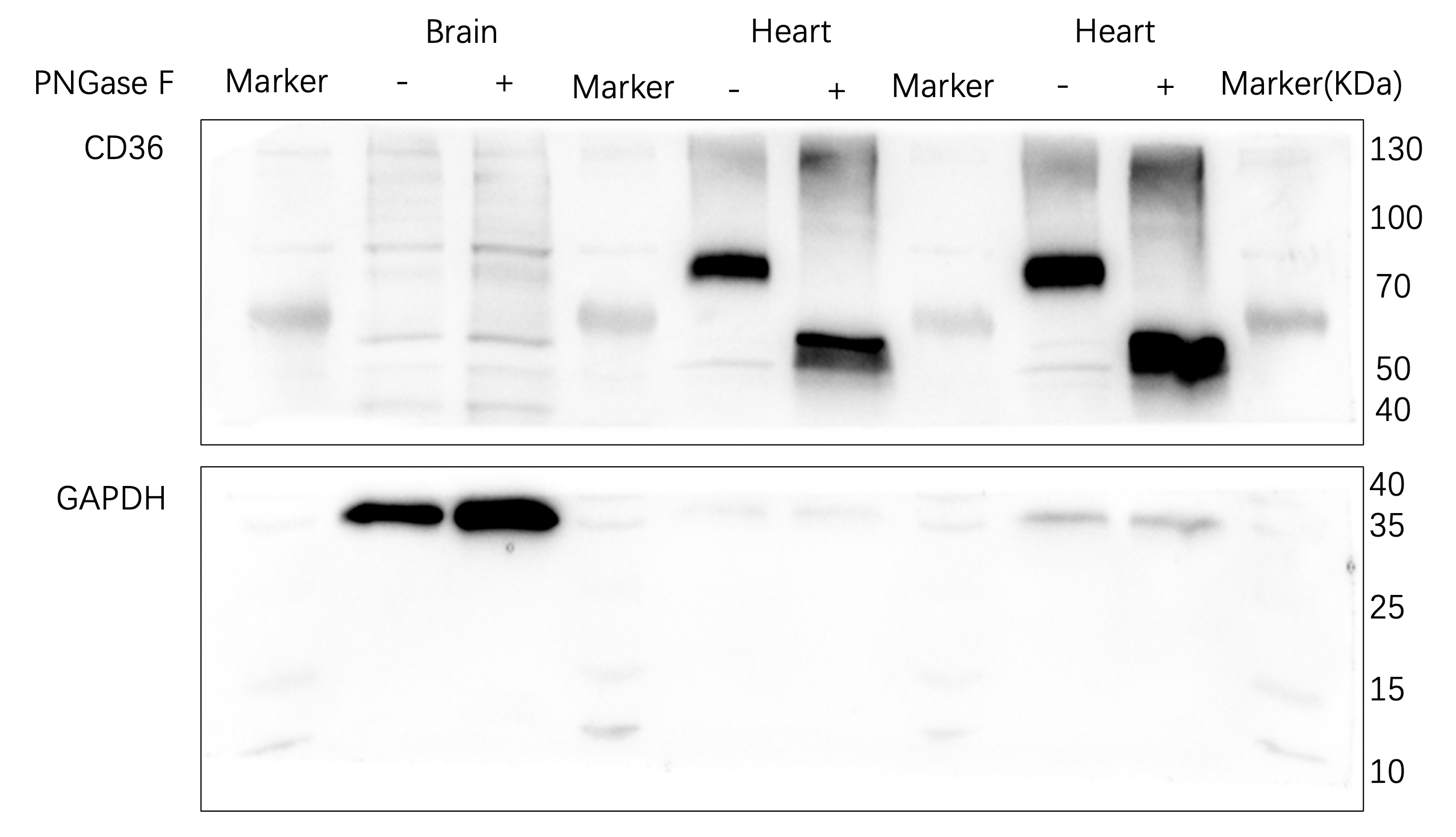

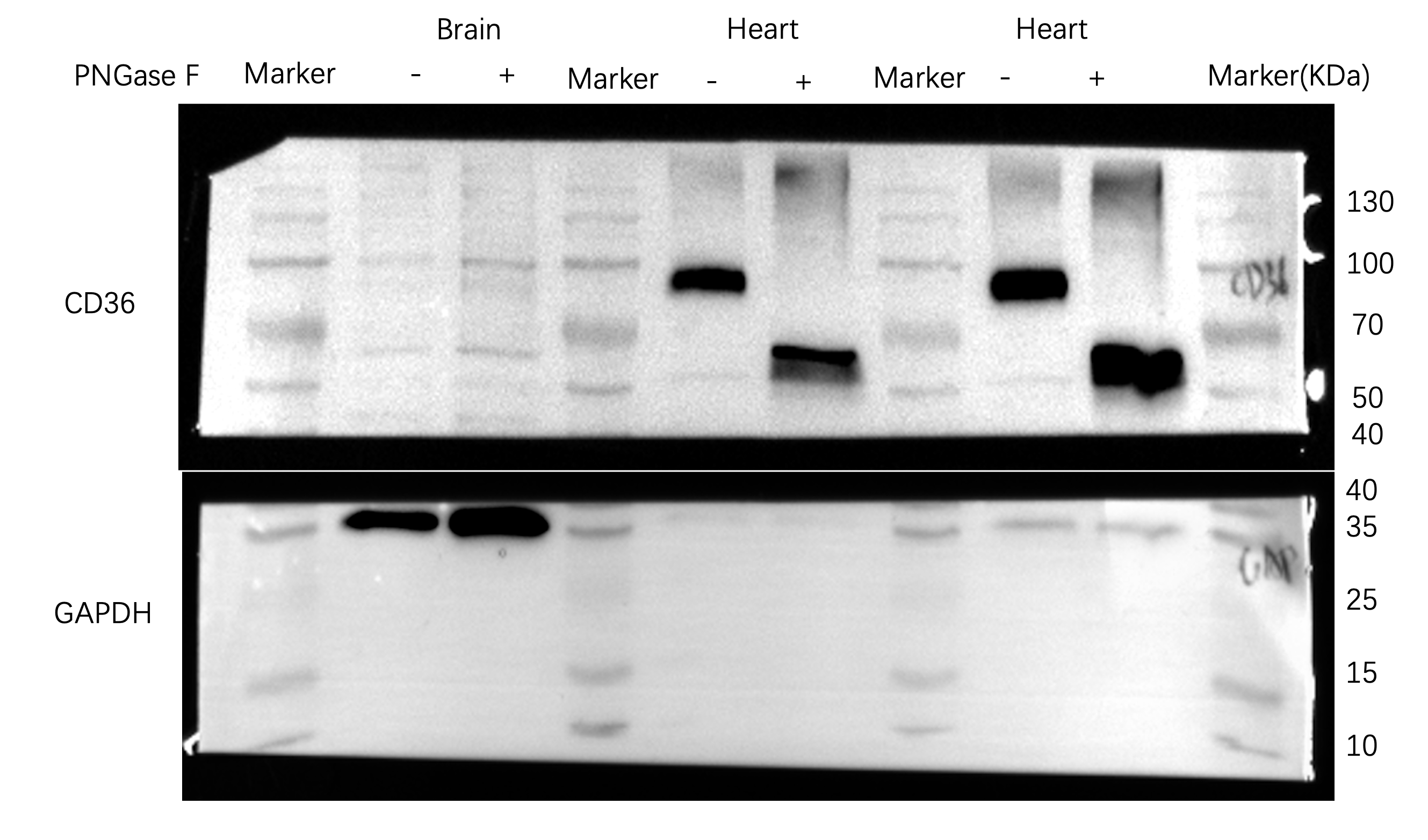

心

-

+

130

100

70

50

35
